# Fine-tuning of SUMOylation modulates drought tolerance

**DOI:** 10.1101/2021.03.29.437542

**Authors:** Xuewei Li, Shuangxi Zhou, Liyuan Lu, Huan Dang, Zeyuan Liu, Huimin Li, Baohua Chu, Pengxiang Chen, Ziqing Ma, Shuang Zhao, Steve van Nocker, Fengwang Ma, Qingmei Guan

## Abstract

SUMOylation is involved in various aspects of plant biology, including drought stress. However, the relationship between SUMOylation and drought stress tolerance is complex; whether SUMOylation has a crosstalk with ubiquitination in response to drought stress remains largely unclear. In this study, we found that both increased and decreased SUMOylation led to increased survival of apple (*Malus* × *domestica*) under drought stress: both transgenic *MdSUMO2A* overexpressing (OE) plants and *MdSUMO2* RNAi plants exhibited enhanced drought tolerance. We further confirmed that MdDREB2A is one of the MdSUMO2 targets. Both transgenic *MdDREB2A* OE and *MdDREB2A*^*K192R*^ OE plants (which lacked the key site of SUMOylation by MdSUMO2A) were more drought tolerant than wild-type plants. However, *MdDREB2A*^*K192R*^ OE plants had a much higher survival rate than *MdDREB2A* OE plants. We further showed SUMOylated MdDREB2A was conjugated with ubiquitin by MdRNF4 under drought stress, thereby triggering its protein degradation. In addition, *MdRNF4* RNAi plants were more tolerant to drought stress. These results revealed the molecular mechanisms that underlie the relationship of SUMOylation with drought tolerance and provided evidence for the tight control of MdDREB2A accumulation under drought stress mediated by SUMOylation and ubiquitination.

## Introduction

Drought stress is one of the main abiotic constraints that limit agricultural development and productivity (Li et al., 2019; Geng et al., 2020). As the global climate warms in the twenty-first century, the frequency of severe drought conditions is increasing (Dai, 2013). Water shortage impairs plant growth and development, limiting plant production and reducing the performance of crop plants and fruit trees (Basu et al., 2016). In fruit trees, water deficit inhibits flower bud differentiation and tree vegetative growth, thereby causing flowers and fruits to drop (Virlet et al., 2015; Niu et al., 2019). To cope with drought stress, plants respond at both the morphological and molecular levels, exhibiting changes in photosynthesis, stomatal movement, hormone content, leaf development, stem extension, root proliferation, hydraulic conductivity, and gene expression (Yordanov et al., 2000; Seiler et al., 2011; Basu et al., 2016; Liao et al., 2016; Sun et al., 2018; Geng et al., 2020; Li et al., 2020). Therefore, decoding the molecular mechanisms that underlie drought responses is critical to the development of new cultivars for future agriculture (Sun et al., 2013b; Liao et al., 2016; Geng et al., 2018; Sun et al., 2018; Li et al., 2020).

Small Ubiquitin-like Modifier (SUMO) is a ∼100-amino-acid polypeptide that is structurally related to ubiquitin (Vierstra and R., 2012). Similar to ubiquitin, SUMOs are encoded as precursor proteins. To attain their mature form, precursor SUMOs require SUMO protease to cleave a C-terminal peptide and expose two consecutive glycine residues that are essential for conjugation to substrates. The biochemical pathway of SUMOylation is also analogous to that of ubiquitination. The first step is SUMO activation, an ATP-dependent reaction that is catalyzed by the heterodimeric E1-activating enzyme, SAE1/SAE2. In the second step, activated SUMO is transferred from SAE to the SUMO-conjugating enzyme (SCE). Finally, the conjugation of SUMO to its substrates is catalyzed by SCE (Colby et al., 2006). The consensus sequence ΨKXE/D (where Ψ is a hydrophobic aliphatic residue; X can be any residue; K, E, and D are standard one-letter symbols for amino acids; and K is the attachment site for SUMO) is considered to be the canonical SUMO attachment site (Novatchkova et al., 2004; Nabil et al., 2014), although other sites may exist. SUMO proteases de-SUMOylate the SUMOylated substrates to recycle SUMO. In addition to its covalent attachment to target proteins, SUMO can also attach to cellular proteins through noncovalent interactions (Galanty et al., 2012).

In addition to the regulation of development and cellular homeostasis under normal growth conditions, SUMOylation is also involved in various biotic and abiotic stress responses, including the response to drought stress (Castro et al., 2012). OsbZIP23 is a SUMOylation substrate that is targeted by the SUMO protease OTS1. SUMOylation of OsbZIP23 causes the transcriptional activation of drought protection genes and improves drought tolerance (Srivastava et al., 2017). Overexpression of SUMO E2-conjugating enzyme (CE) in rice (*Oryza sativa*) impairs drought tolerance by reducing the accumulated proline content relative to the wild type (Nurdiani et al., 2018). However, overexpression of SaSce9 from the halophyte grass *Spartina alterniflora* enhances salinity and drought stress tolerance in Arabidopsis (Karan and Subudhi, 2012), indicating that CE plays various roles in different plants. The SUMO E3 ligase MMS21 negatively influences Arabidopsis drought response through an ABA-dependent pathway (Zhang et al., 2013). Likewise, the rice SUMO protease OsOTS1 also plays a negative role in drought stress response through an ABA-dependent pathway (Srivastava et al., 2017). Another SUMO E3 ligase, SIZ1, plays more complicated roles in drought stress response in different plant species. Rice Os*SIZ1* confers drought tolerance in transgenic bentgrass and cotton (Neelam et al., 2017). Transgenic tobacco plants ectopically expressing tomato *SlSIZ1* are more tolerant to drought stress (Zhang et al., 2017b), as are Arabidopsis plants overexpressing *SIZ1*(Zhang et al., 2013). However, *siz1* mutant plants displayed drought-sensitive or drought-tolerant phenotypes in three independent studies (Catala et al., 2007; Miura et al., 2013; Kim et al., 2017). Reports on Arabidopsis SIZ1 overexpressing (OE) plants and *siz1* mutants indicate that either increased or decreased SUMOylation levels can improve drought resistance. However, the physiological and molecular basis for this effect is unclear. In addition, despite the identification of drought-related SUMO targets, the biological function of their SUMOylation modification is largely unknown.

The SUMO-interacting proteins (SIPs) play a crucial role in the regulation of SUMOylated proteins; they usually interact with SUMO through SUMO-interacting motifs (SIMs). Proteins with SIMs include a group of RING-type ubiquitin E3 ligases, DNA methyltransferses or demethylases, and histone methyltransferses or demethylases (Nabil et al., 2014; Kumar et al., 2017). The best-studied SIPs are the RING-type ubiquitin E3 ligases that target SUMOylated proteins for degradation by the proteasome pathway. RING finger protein 4 (RNF4, also known as SUMO-targeted ubiquitin E3 ligase or STUbL) ubiquitinates promyelocytic leukemia protein (PML) or the nuclear receptor NR4A1 that has been SUMOylated by SUMO2/3 in mammals (Valérie et al., 2008; Geoffroy and Hay, 2009; Zhang et al., 2017a). RNF4 also ubiquitinates SUMOylated proteins in the fission yeast *Schizosaccharomyces pombe* (Sun et al., 2007) and promotes the ubiquitination of activated MEK1 in a RING-finger-dependent manner in *Dictyostelium*(Sobko et al.). SUMOylation of PML recruits RNF4, triggering its Lys 48-linked polyubiquitination and degradation (Tatham et al., 2008; Valérie et al., 2008). NR4A1 is SUMOylated by SUMO2/3 and targeted by RNF4 for polyubiquitination and subsequent degradation to control macrophage cell death (Zhang et al., 2017a). Although several studies have reported the important and conserved role of RNF4 in multicellular eukaryotes, only one study has investigated the role of AT-STUbL4 in the floral transition in plants (Nabil et al., 2014) *at-stubl4* mutant plants flowered later than the wild type, whereas AT-STUbL4 OE plants flowered earlier (Nabil et al., 2014). To date, it remains unclear whether RNF4 can recognize and ubiquitinate SUMOylated proteins in plants, especially during the response to drought stress.

Dehydration-responsive element-binding factor (DREB2A) is a transcription factor that binds specifically to the DRE/CRT *cis*-element and is rapidly induced by dehydration (Liu et al., 1998; Li et al., 2019). DREB2A is a key factor in plant drought stress tolerance. Overexpression of full-length DREB2A in apple, *Pennisetum glaucum, Zea mays*, and *O. sativa* enhances tolerance to drought stress (Agarwal et al., 2007; Qin et al., 2007; Cui et al., 2011; Liao et al., 2016). In Arabidopsis, DREB2A is unstable under control conditions owing to its negative regulatory domain (NRD) (Sakuma et al., 2006b; Qin et al., 2008; Mizoi et al., 2013; Sadhukhan et al., 2014; Morimoto et al., 2017). The overexpression of *DREB2A-CA* (a constitutively active form of DREB2A with the NRD domain deleted) increases drought tolerance in Arabidopsis (Sakuma et al., 2006b). Various posttranslational modifications of DREB2A, including SUMOylation and ubiquitination, are tightly associated with its stability and transcriptional activity in Arabidopsis (Qin et al., 2008; Wang et al., 2020). Two types of ubiquitin E3 ligase, BPMs and DRIPs, mediate the degradation of DREB2A in Arabidopsis (Qin et al., 2008; Morimoto et al., 2017). However, SUMOylation of DREB2A by SCE1 can repress the interaction between DREB2A and BPM2, thereby increasing DREB2A protein stability under high temperature (Wang et al., 2020). Whether SUMOylated DREB2A can be targeted by ubiquitin E3 ligases for degradation remains unclear.

Apple DREB2A does not contain the NRD domain that is targeted for degradation in Arabidopsis. Unlike Arabidopsis DREB2A, MdDREB2A is consistently stable under normal conditions (Li et al., 2019). Overexpression of *MsDREB6*.*2* (a *MdDREB2A* homolog) or *MpDREB2A* enhanced drought tolerance of apple or Arabidopsis (Liao et al., 2016; Li et al., 2019). Given the lack of an NRD domain in MdDREB2A, its posttranslational modifications and the molecular mechanisms of its protein stability under stress require clarification. In this study, we found that both increased and decreased SUMOylation levels improved apple drought tolerance. We further identified MdDREB2A as one of the SUMOylation target proteins and demonstrated that SUMOylation of MdDREB2A was critical for protein stability and drought tolerance. In addition, we further provided evidence that SUMOylated MdDREB2A could be recognized and ubiquitinated by MdRNF4 under drought stress, leading to the degradation of MdDREB2A. Our results highlight the roles of SUMOylation in apple drought tolerance and provide insight into the RNF4-mediated ubiquitination of SUMOylated MdDREB2A in response to drought.

## Results

### Expression patterns and localization of SUMO2s in apple

The apple genome contains six SUMO2 genes (Fig. 1A). Due to genome duplication, each pair of genes on different chromosomes has almost identical coding sequences, and we therefore named the three pairs MdSUMO2A, MdSUMO2B, and MdSUMO2C. Protein alignment revealed a protein sequence similarity of 76%–88% among these three MdSUMO2 proteins (Fig. 1A). To characterize the function of apple SUMO2 proteins in response to drought stress, we first examined their expression patterns under drought. We found that the *MdSUMO2*s had similar expression patterns in response to drought (Fig. 1B), suggesting that they may have similar functions under drought stress. Apple *SUMO2A* and *SUMO2B* were more abundant in all tissues, whereas *SUMO2C* was less abundant in all tissues examined (Fig. S1A). When *MdSUMO2A::GUS* was ectopically expressed in Arabidopsis, similar results were observed, and GUS signal was detected in all tissues (Fig. S1B– I).

**Fig. 1.**
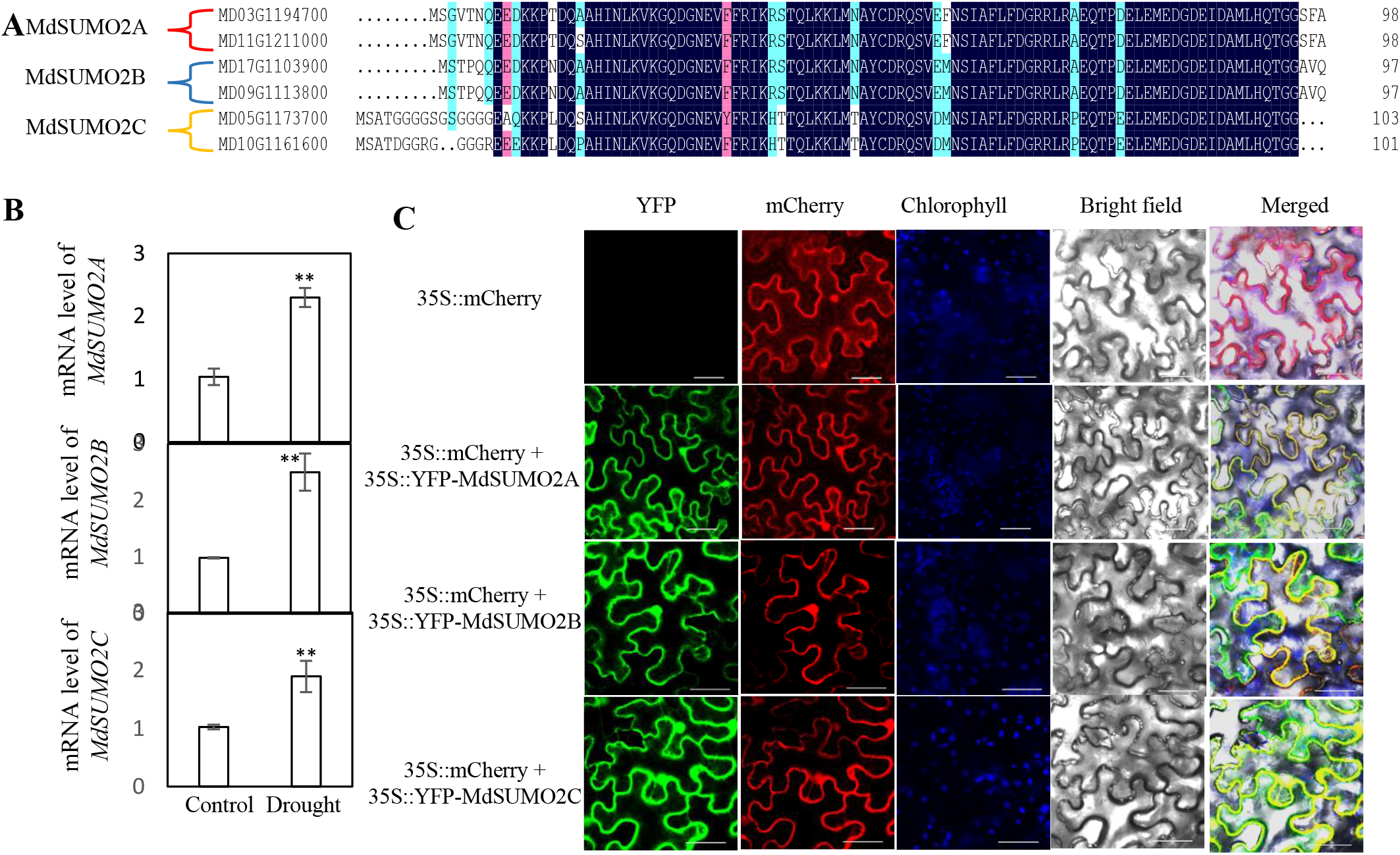
Sequences, responses to drought stress, and localization of MdSUMO2s. **(A)** Comparison of amino acid sequences of MdSUMO2A, MdSUMO2B, MdSUMO2C in apple. MD03G1194700 and MD11G1211000 were named MdSUMO2A; MD17G1103900 and MD09G1113800 were named MdSUMO2B; MD5G1173700 and MD10G1161600 were named MdSUMO2C. **(B)** *MdSUMO2* expression in response to drought in 2-month old GL-3 plants which were exposed to drought for 0 and 6 days. **(C)** Subcellular localization of MdSUMO2s. YFP-MdSUMO2A, YFP-MdSUMO2B, or YFP-MdSUMO2C was transformed into 5-week-old tobacco (*Nicotiana benthamiana*) leaves for 3 days, and YFP and mCherry fluorescent signals were then observed. Bars = 40 μm. Error bars indicate standard error (n = 3). Asterisks indicate significant differences based on one-way ANOVA and Tukey test (**, *P* < 0.01).

We aligned SUMO2A proteins from different plant species and found that their sequences were highly conserved throughout the plant kingdom (Fig. S2A). MdSUMO2A was highly similar to SUMO2A from *Prunus mume* (Fig. S2B). We then cloned SUMO2A, SUMO2B, and SUMO2C from the apple genome. Co-localization with mCherry suggested that apple SUMO2A, SUMO2B, and SUMO2C are localized in the nucleus, plasma membrane, and cytoplasm (Fig. 1C).

### Knocking down *MdSUMO2s* or knocking in one *MdSUMO2* gene leads to drought stress tolerance

To understand the biological function of the MdSUMO2s, we generated a series of transgenic plants: *MdSUMO2A* OE (over expression) with a higher *MdSUMO2A* expression level; *MdSUMO2A* RNAi with reduced expression of *MdSUMO2A* only; and *MdSUMO2* RNAi with reduced expression of *MdSUMO2A, MdSUMO2B*, and *MdSUMO2C* (Fig. S3).

After transplant, the transgenic plants and non-transgenic plants (GL-3) were exposed to prolonged drought stress by maintaining soil volumetric water content of 18-23% for three months. As shown in Fig. 2A and B, long-term moderate drought stress reduced the growth of all plants. However, compared with GL-3 plants, *MdSUMO2A* OE plants were taller, and *MdSUMO2* RNAi plants were shorter (Fig. 2A-B, Fig. S4). In addition to differences in plant height, *MdSUMO2A* OE plants had greater stem diameters and longer internodes than GL-3 plants under drought stress, whereas *MdSUMO2* RNAi plants had smaller stem diameters and shorter internodes (Fig. S5). Moreover, *MdSUMO2A* OE plants had greater shoot dry weights under control and drought conditions, whereas *MdSUMO2* RNAi plants had lower aboveground biomass (Fig. S6). These results indicate that *MdSUMO2A* OE plants grew more vigorously under long-term drought, whereas *MdSUMO2* RNAi plants grew more slowly.

**Fig. 2.**
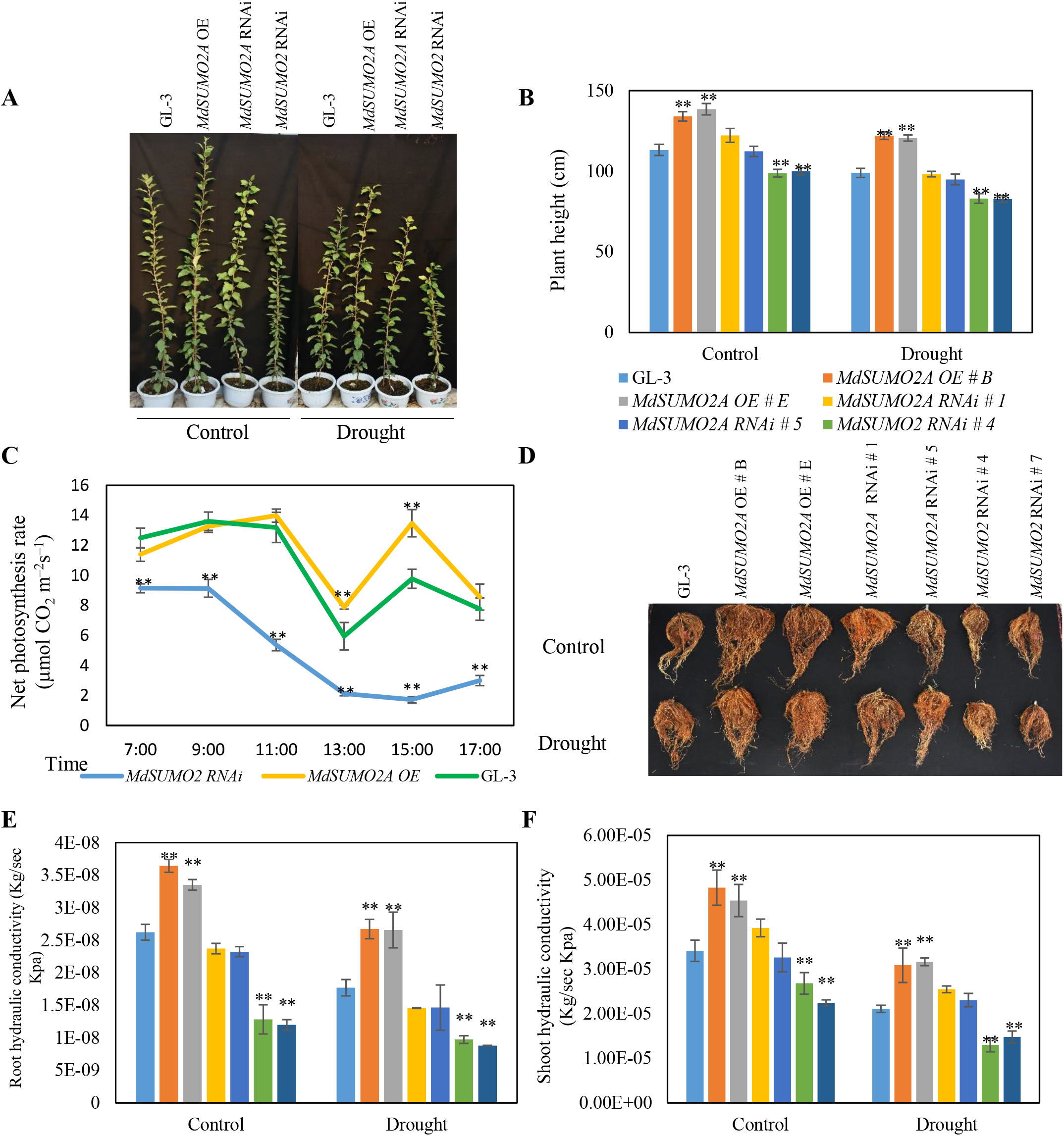
*MdSUMO2A* OE plants show increased tolerance to drought stress. **(A)** Morphology of *MdSUMO2* transgenic plants under control and long-term drought stress. **(B)**-**(F)** Plant height **(B)**, net photosynthesis **(C)**, root morphology **(D)**, root hydraulic conductivity **(E)**, and shoot hydraulic conductivity **(F)** of GL-3 and *MdSUMO2* transgenic plants under control and long-term drought stress. Plants were exposed to drought for up to 3 months. During treatment, 43-48% soil volumetric water content (VWC) was maintained as control and 18-23% of VWC was maintained as drought treatment. Error bars indicate standard error [n = 12 in (B), 7 in (C), 5 in (E) and (F)]. Asterisks indicate significant differences based on one-way ANOVA and Tukey test (*, *P* < 0.05; **, *P* < 0.01). OE, overexpression.

Drought can adversely affect crop photosynthetic capacity, water use efficiency, and yield (Xu et al., 2008; Sun et al., 2013a; Mao et al., 2015), and drought stress reduced the photosynthetic capacity of all plants in the current experiment (Fig. S7). However, *MdSUMO2A* OE plants had a greater photosynthetic capacity than GL-3 plants under drought stress, whereas that of *MdSUMO2A* RNAi plants was lower (Fig. S7A). Under drought stress, stomatal conductance and transpiration rate were also higher in *MdSUMO2A* OE plants than in GL-3 plants, and both parameters were lower in *MdSUMO2* RNAi plants (Fig. S7B-C). We also measured the photosynthetic capacity of GL-3 and transgenic plants under drought during the daytime from 7:00 AM to 5:00 PM. Similar results were observed. That is, *MdSUMO2A* OE plants maintained the highest photosynthetic rate under drought stress and exhibited a higher transpiration rate and stomatal conductance after noon, whereas *MdSUMO2* RNAi plants had the lowest values for these parameters (Fig. 2C and Fig. S8).

The root system plays an important role in plant drought resistance (Liao et al., 2016; Geng et al., 2018; Hu et al., 2018). After long-term drought, the root systems of *MdSUMO2A* OE plants were much more extensive (Fig. 2D), as indicated by root dry weight in Fig. S10. However, the root systems of *MdSUMO2* RNAi plants were much smaller than those of GL-3 under control and drought conditions (Fig. 2D and Fig. S9). Consistent with their strong root systems and greater shoot growth, *MdSUMO2A* OE plants had higher hydraulic conductivity of roots and shoots (Fig. 2E and F), whereas those of *MdSUMO2* RNAi plants were lower. These results suggest that *MdSUMO2A* OE plants performed better under drought, exhibiting vigorous shoot and root growth, as well as higher hydraulic conductivity and photosynthetic capacity.

Leaf morphology is important for drought tolerance (Anyia and Herzog, 2004; Sun et al., 2013a; Wu et al., 2014). *MdSUMO2* RNAi leaves were smaller than those of GL-3 and *MdSUMO2A* OE plants under control and drought conditions (Fig. 3A), as indicated by leaf area measurements in Fig. 3B. Leaf lengths and widths were also smaller in *MdSUMO2* RNAi plants under control and drought conditions (Fig. S10A-B). Likewise, under both conditions, single-leaf dry weight was much lower in *MdSUMO2* RNAi plants than in GL-3 and *MdSUMO2A* OE plants (Fig. S10C). *MdSUMO2* RNAi leaves were much thicker than GL-3 and *MdSUMO2A* OE leaves under control and drought conditions (Fig. 3C and D). Consistently, *MdSUMO2* RNAi leaves had a greater water holding capacity (Fig. 3E). By contrast, the leaf area, dry weight, thickness, and water holding capacity of *MdSUMO2A* OE leaves were comparable to those of GL-3 leaves under control and drought conditions (Fig. 3A– E). Water use efficiency was measured using ^13^C, and *MdSUMO2* RNAi plants maintained a higher WUE than GL-3 and *MdSUMO2* OE plants under control and drought conditions (Fig. 3F). Plants accumulate the phytohormone abscisic acid (ABA) after drought stimulus (Zhu, 2016). After drought stress, the ABA content of *MdSUMO2A* OE plants was lower than that of GL-3 plants, whereas that of *MdSUMO2* RNAi plants was higher (Fig. S11). These results suggest that *MdSUMO2* RNAi plants resist drought by adjusting their leaf morphology, increasing their WUE, and accumulating more ABA.

**Fig. 3.**
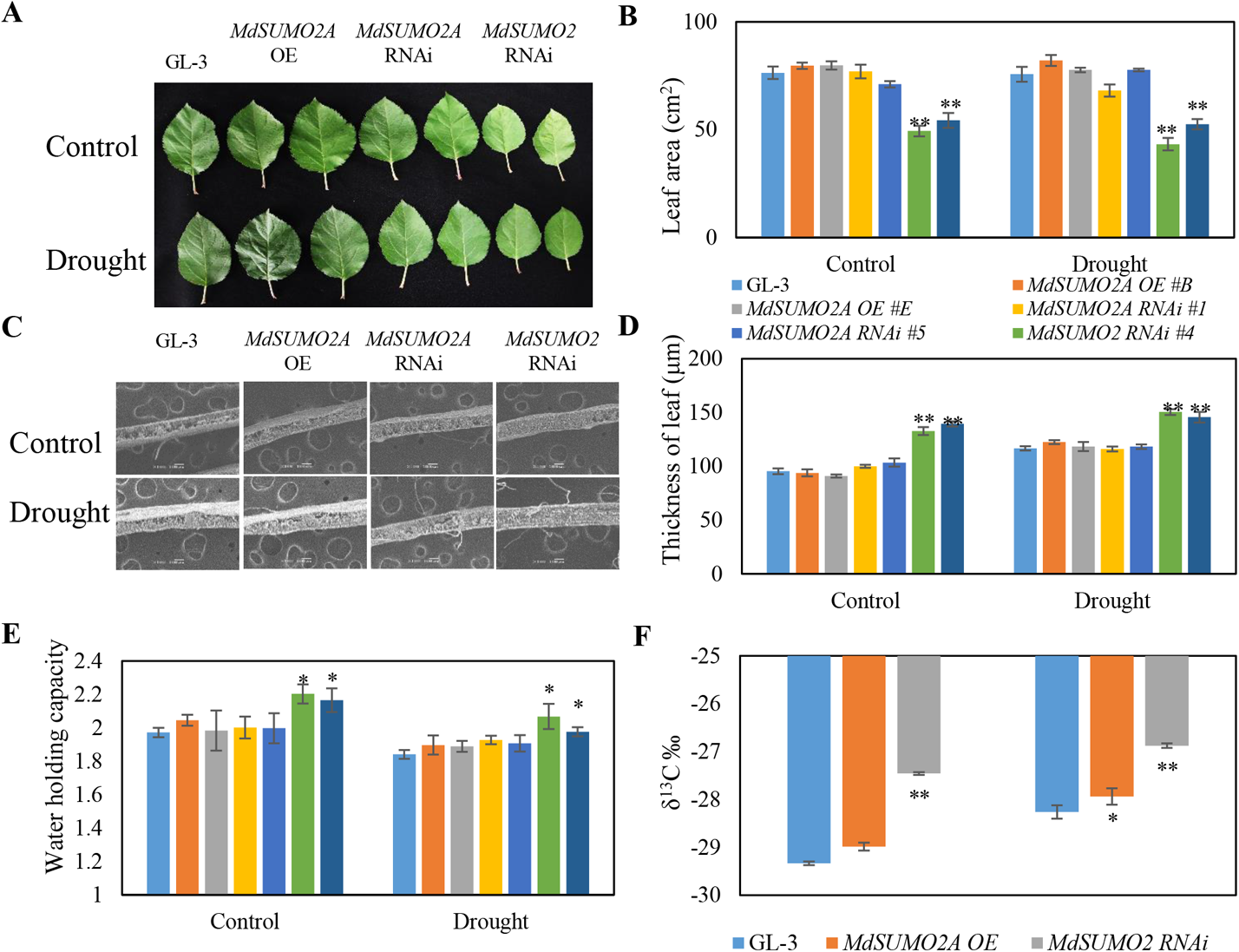
*MdSUMO2* RNAi plants display increased tolerance to drought stress. **(A)-(F)** Leaf morphology **(A)**, leaf area **(B)**, leaf thickness **(C** and **D)**, water holding capacity **(E)**, and water use efficiency **(F)** of GL-3 and *MdSUMO2* transgenic plants under control and long-term drought stress. Leaf thickness was observed using tungsten filament scanning electron microscope (TEM); water holding capacity = (leaf saturated weight-dry weight) /dry weight; water use efficiency was detected by carbon isotope (^13^C) composition. Plants were exposed to drought for up to 3 months. During treatment, 43-48% soil volumetric water content (VWC) was maintained as control and 18-23% of VWC was maintained as drought treatment. Error bars indicate standard error [n = 9 in (B), 16 in (D), 6 in (E), 3 in (F)]. Asterisks indicate significant differences based on one-way ANOVA and Tukey test (*, *P* < 0.05; **, *P* < 0.01). OE, overexpression.

There were no significant differences in the parameters mentioned above between *MdSUMO2A* RNAi and GL-3 plants under drought stress (Fig. 1–4, Fig. S4–12), suggesting that the MdSUMO2s have redundant functions in response to drought.

**Fig. 4.**
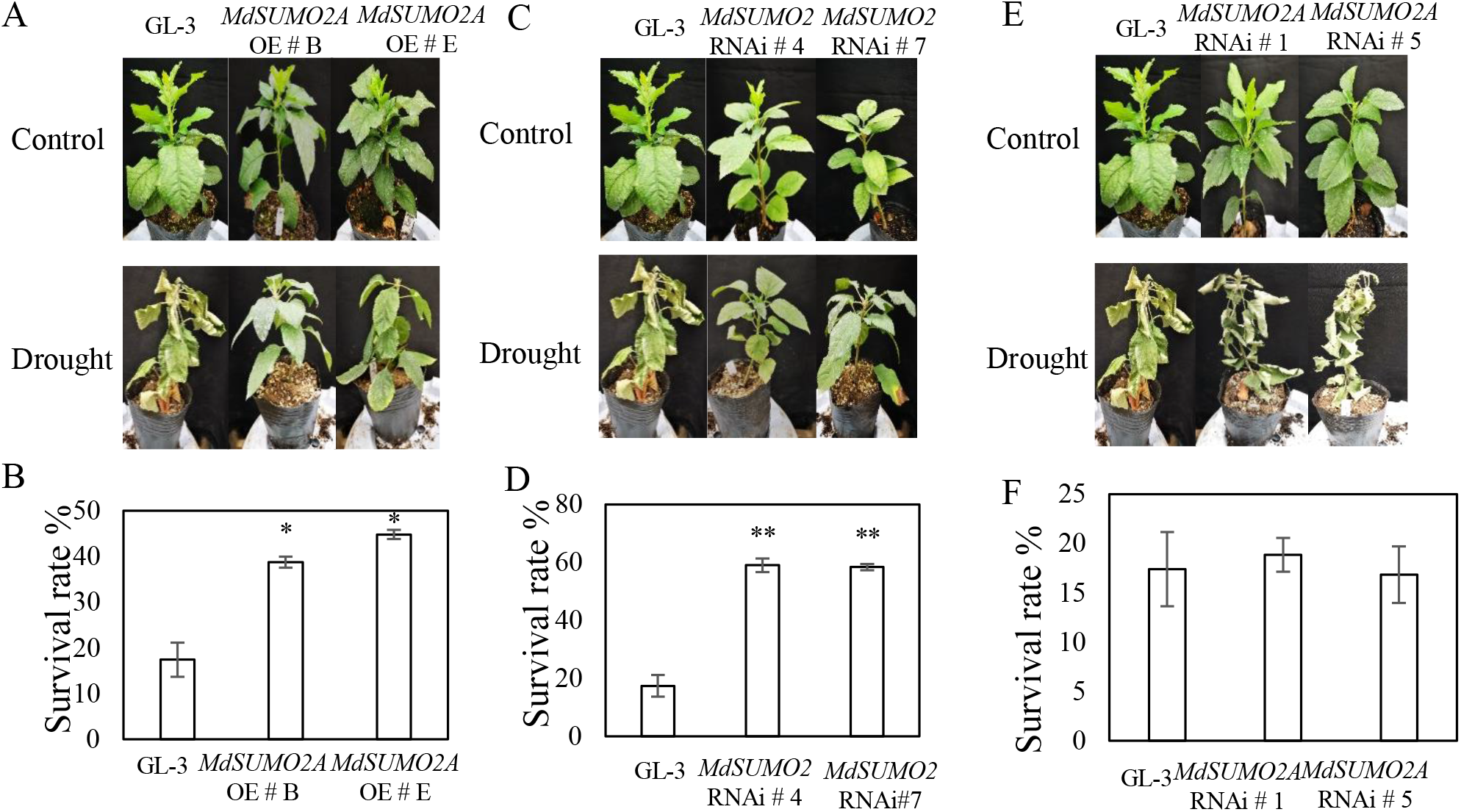
Tolerance of *MdSUMO2* transgenic plants and GL-3 in response to short-term drought stress. **(A)** Tolerance of *MdSUMO2A* OE and GL-3 plants under short-term drought. **(B)** Survival rate of plans shown in **(A). (C)** Tolerance of *MdSUMO2* RNAi and GL-3 plants under short-term drought. **(D)** Survival rate of plans shown in **(C). (E)** Tolerance of *MdSUMO2A* RNAi and GL-3 plants under short-term drought. **(F)** Survival rate of plans shown in **(E)**. Error bars indicate standard error (n = 3). Asterisks indicate significant differences based on one-way ANOVA and Tukey test (*, *P* < 0.05; **, *P* < 0.01). OE, overexpression.

To further support the notion that both *MdSUMO2A* OE and *MdSUMO2* RNAi plants were tolerant to drought stress, we treated all plants with a shorter-term drought stress. After 3 weeks of drought treatment, 83% of the GL-3 plants had wilted, whereas 40% of the *MdSUMO2A* OE plants and 58% of the *MdSUMO2* RNAi plants were still alive (Fig. 4A-D), suggesting that the *MdSUMO2* RNAi plants had a higher survival capacity than the *MdSUMO2A* OE plants. By contrast, *MdSUMO2A* RNAi plants did not differ in survival rate from GL-3 plants under drought stress (Fig. S4E-F). We also performed an extreme drought treatment after the long-term drought treatment by withholding water for 10 days. Both the *MdSUMO2* RNAi and *MdSUMO2A* OE plants performed better than the GL-3 plants under drought, and the *MdSUMO2* RNAi plants were more drought tolerant than the *MdSUMO2A* OE plants (Fig. S4B). All these data suggest that *MdSUMO2A* OE and *MdSUMO2* RNAi plants were more drought tolerant than GL-3 plants and that *MdSUMO2* RNAi plants had higher survival ability than *MdSUMO2A* OE plants.

In addition, we examined the SUMOylation of GL-3 and *MdSUMO2* transgenic plants under control and prolonged drought stress conditions. As shown in Fig. S12, *MdSUMO2A* OE plants had a slightly higher SUMOylation level than GL-3 plants under control and drought conditions, whereas the SUMOylation level of *MdSUMO2* RNAi plants was lower.

### Identification of MdSUMO2 targets reveals SUMOylation of MdDREB2A by MdSUMO2s

To identify potential targets of MdSUMO2 proteins, we performed proteomic analysis according to previous methods (Miller et al., 2010; Miller and Vierstra, 2011). Since Arabidopsis SUMO1 has high sequence similarity with MdSUMO2 (Fig. S3A), we used the anti-SUMO1 antibody to recognize three MdSUMO2s. After mass spectrometry, we identified 1314 potential targets of MdSUMO2A (Supplemental Data Set 1), including MdDREB2A, MdALI, MdAQP2, MdHSP20, MdH2B, MdCAT2, and MdbZIP (Fig. S13). Using a SUMOylation reconstitution assay in *Escherichia coli* in which MdSUMO2 and the candidate substrates were expressed (Elrouby and Coupland, 2010), we verified the SUMOylation of MdDREB2A, MdAQP2, and MdALI by the MdSUMO2s (Fig. 5A-G). Three and one lysine sites are potential SUMO conjugation sites in MdDREB2A and MdAQP2, respectively. To determine the actual SUMOylation sites, each candidate lysine (K) was replaced by arginine (R) singly or in combinations. SUMOylation assays using the *E. coli* system suggested that K192 and K272 were required for MdSUMO2A-mediated SUMOylation of MdDREB2A and MdAQP2, respectively (Fig. 5A and E). In addition, K192 was also required for MdDREB2A SUMOylation by MdSUMO2C, whereas K192, K217, and K369 were required for MdDREB2A SUMOylation by MdSUMO2B (Fig. 5B and C). For MdALI, there are five lysine sites and one SIM for potential SUMO conjugation. Deleting the SIM or mutating each lysine to R could not abolish the SUMOylation of MdALI by MdSUMO2A (Fig. S14). However, mutation of all five lysine sites to R or mutation of four lysine sites to R and in combination with SIM deletion could almost completely abolish the SUMO conjugation by MdSUMO2A, indicating that these five lysine sites and the SIM were all required for SUMOylation of MdALI by MdSUMO2A (Fig. 5G).

**Fig. 5.**
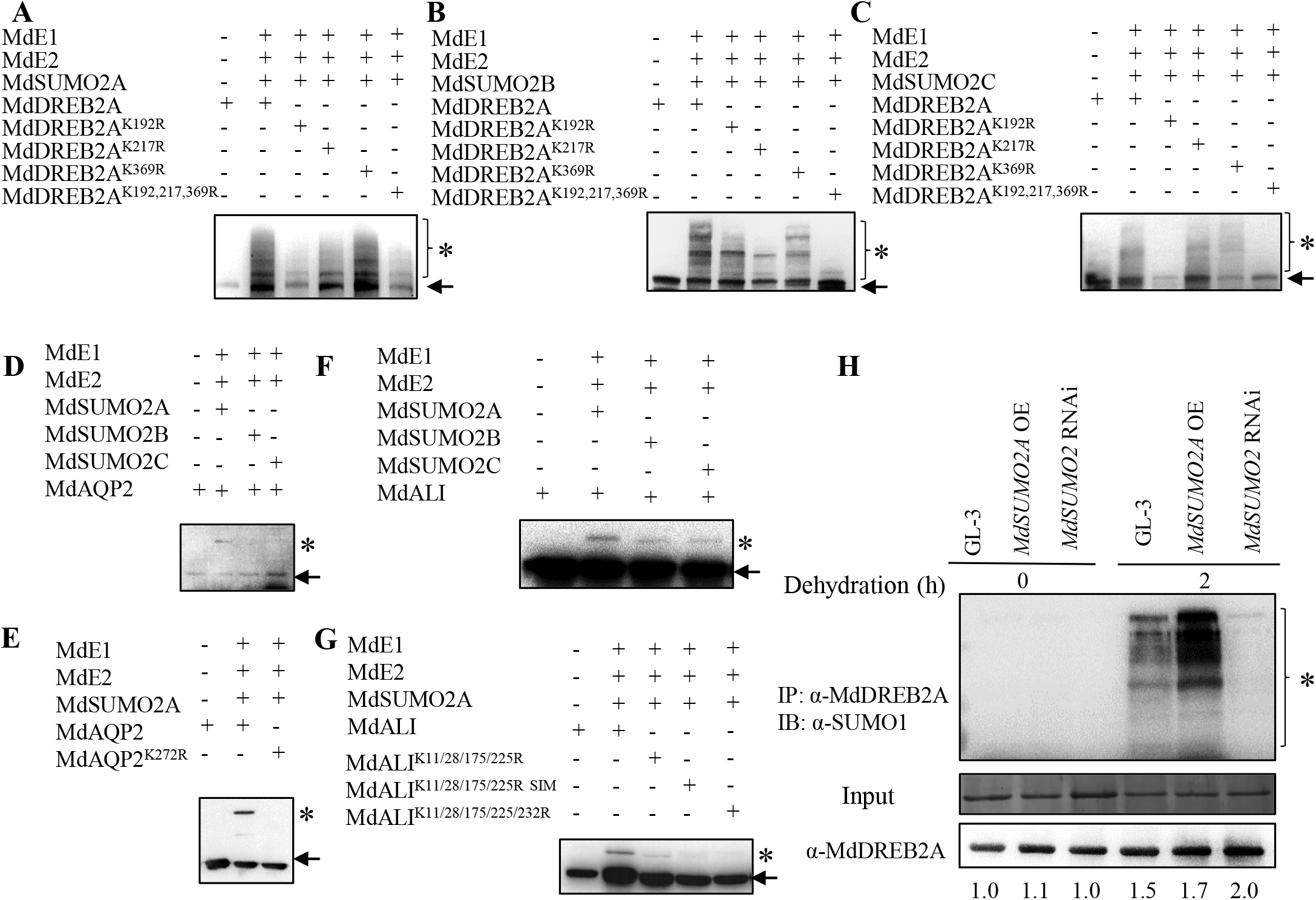
SUMOylation of MdDREB2A, MdAQP2, and MdALI using *E. Coli* system. **(A)-(C)** MdDREB2A was SUMOylated by MdSUMO2A, MdSUMO2B, and MdSUMO2C. Putative SUMOylation sites (K) of MdDREB2A were mutated to arginine (R). **(D)** and **(E)** SUMOylation of AQP2 by MdSUMO2A. Putative SUMOylation site (K272) of MdAQP2 was mutated to arginine (R). **(F)** and **(G)** SUMOylation of MdALI by MdSUMO2A, MdSUMO2B, and MdSUMO2C. Putative SUMOylation sites (K) or SIM of MdALI was mutated to arginine (R). **(H)** SUMOylation of MdDREB2A and MdDREB2A protein in GL-3, *MdSUMO2* RNAi and *MdSUMO2A* OE plants under control or dehydration conditions. * indicates SUMOylated substrates; arrows indicate substrates. OE, overexpression.

Because it is an important factor in plant drought stress response (Sakuma et al., 2006b; Chen et al., 2007; Qin et al., 2007; Reis et al., 2014), we next focused on MdDREB2A. Since MdDREB2A could be SUMOylated in the *E*.*Coli* system, and MdDREB2A did not contain the SIM, we tested the interaction of MdDREB2A and MdCE, the SUMO E2-conjugating enzyme. MST and CO-IP analysis revealed that MdCE interacts with MdDREB2A *in vitro and vivo* (Fig. S15). SUMOylation can affect target protein localization, protein–protein interaction, and protein stability. We co-localized MdSUMO2A with MdDREB2A and found that SUMOylation of MdDREB2A did not affect its subcellular localization (Fig. S16). We also examined the effect of SUMOylation on the stability of MdDREB2A. As shown in Fig. 5H, MdDREB2A protein level was significantly increased in the *MdSUMO2* RNAi plants under drought conditions but also slightly higher in the *MdSUMO2A* OE plants.

In addition to the *in vitro* SUMOylation of MdDREB2A, we also examined the *in vivo* SUMOylation of MdDREB2A by MdSUMO2 under control and drought conditions. After immunoprecipitation using anti-MdDREB2A antibody, SUMOylation of MdDREB2A was detected in GL-3 plants under drought stress, but much less SUMOylation was observed in *MdSUMO2* RNAi plants (Fig. 5H).

### SUMOylation of MdDREB2A is critical for drought stress tolerance and is coupled with ubiquitination during drought

DREB2A is a positive regulator of plant drought and heat stress tolerance (Kim et al., 2011; Meng et al., 2011; Li et al., 2019). Arabidopsis wild-type plants overexpressing *DREB2A*^*K163R*^ (in which K was mutated to R) exhibited decreased thermotolerance (Wang et al., 2020). We therefore examined whether SUMOylation of MdDREB2A affected apple drought stress resistance. We transformed 35S::*MdDREB2A* (*MdDREB2A* OE) and 35S::*MdDREB2A*^*K192R*^ (*MdDREB2A*^*K192R*^ OE, in which K192 was mutated to arginine) into wild-type GL-3 apple plants. Both transgenic plants had better survival ability under drought stress compared with the wild type (Fig. 6A-B). However, *MdDREB2A*^*K192R*^ OE plants had a higher survival rate than *MdDREB2A* OE plants (Fig. 6A-B). In addition, after drought stress, *MdDREB2A* OE plants had higher photosynthetic capacity than *MdDREB2A*^*K192R*^ OE plants (Fig. 6C). These data suggest that SUMOylation of MdDREB2A tightly controls plant drought tolerance.

**Fig. 6.**
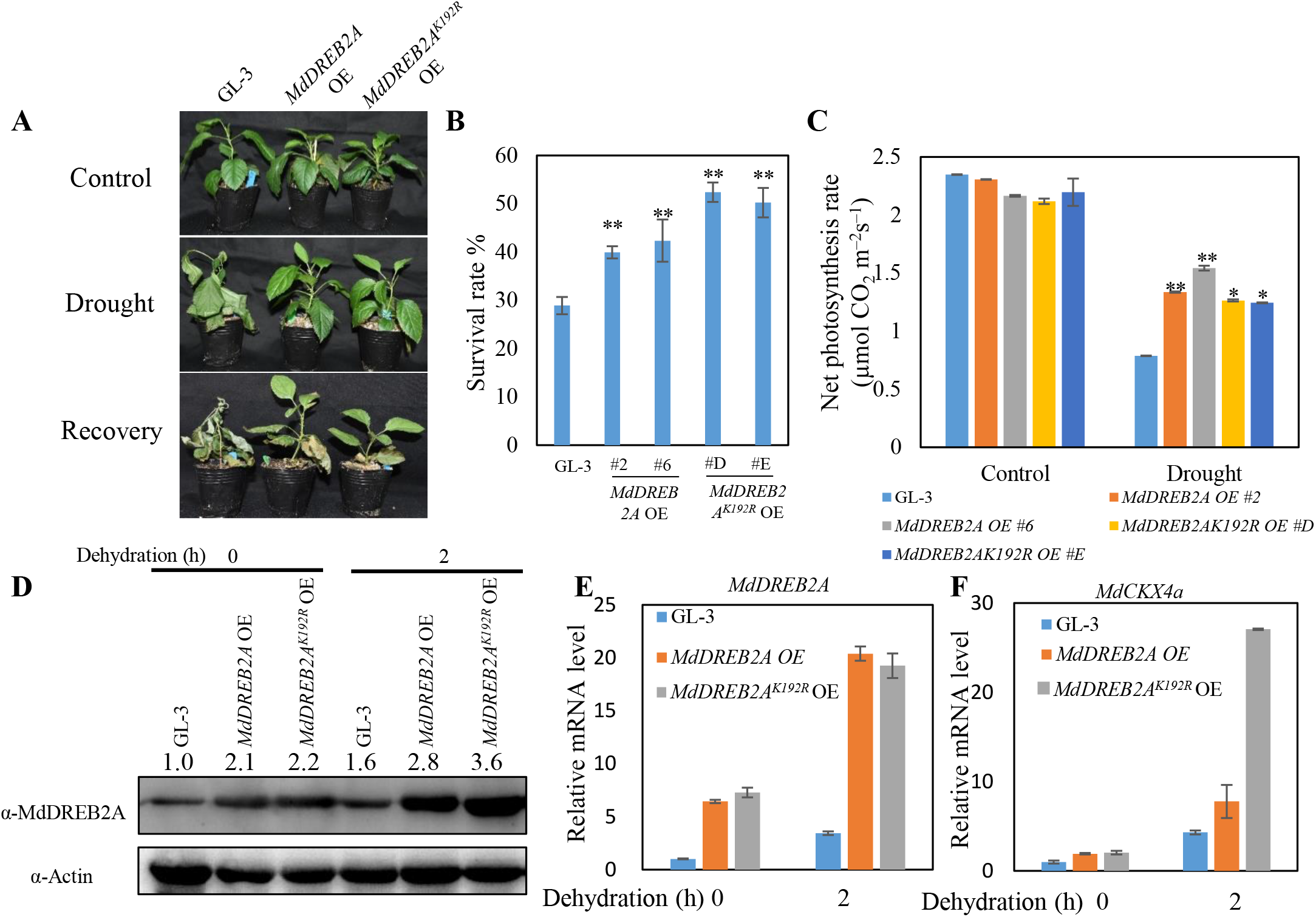
SUMOylation of MdDREB2A is critical for drought stress tolerance. **(A)** Morphology of *MdDREB2A* OE and *MdDREB2A*^*K192R*^ OE transgenic plants under drought treatment for 3 weeks. **(B)** Survival rate of the plants shown in **(A). (C)** Net photosynthesis rate of *MdDREB2A* OE and *MdDREB2A*^*K192R*^ OE transgenic plants under drought treatment. **(D)** MdDREB2A accumulation in *MdDREB2A*^*K192R*^ OE and *MdDREB2A* OE plants under dehydration treatment. **(E)** and **(F)** mRNA level of *MdDREB2A* and *MdCKX4a* in *MdDREB2A* OE and *MdDREB2A*^*K192R*^ OE transgenic plants under dehydration treatment. Error bars indicate standard error [n = 4 in (B), 13 in (C), 3 in (E) and (F)]. Asterisks indicate significant differences based on one-way ANOVA and Tukey test (*, *P* < 0.05; **, *P* < 0.01). OE, overexpression.

Because SUMOylation can affect protein stability, we then examined MdDREB2A protein levels in both transgenic plants under control and drought conditions. As shown in Fig. 6D, both transgenic plants had more MdDREB2A than GL-3 plants under control conditions. Under drought conditions, *MdDREB2A* OE plants accumulated more MdDREB2A protein than GL-3 plants, but less than transgenic plants carrying 35S::*MdDREB2A*^*K192R*^ (Fig. 6D). In addition, the transcripts of MdDREB2A were comparable between two transgenic plants (Fig. 6E). We also transformed 35S::*MdDREB2A* and 35S::*MdDREB2A*^*K192R*^ into apple calli and found that transgenic calli carrying either constructs were more tolerant to simulated drought treatment than wild-type calli. Furthermore, calli carrying 35S::*MdDREB2A*^*K192R*^ were more tolerant to PEG than calli carrying 35S::*MdDREB2A* (Fig. S17). In apple, MdDREB2A targets *MdCKX4a* to modulate drought tolerance (Liao et al., 2016). We next evaluated the *MdCKX4a* expression in transgenic plants and GL-3. As shown in Fig. 6F, *MdCKX4a* expression was higher in transgenic apple plants under normal and drought conditions and much higher in plants expressing 35S::*MdDREB2A*^*K192R*^ than in plants carrying 35S::*MdDREB2A*. These results indicate that SUMOylation of MdDREB2A is important for its stability and activity.

The above phenomena prompted us to investigate whether other protein modifications were involved. Indeed, we found that MdDREB2A accumulation was similar in both genotypes of transgenic plants under drought stress when they were treated with MG132, a 26S proteasome inhibitor (Fig. 7A). The 26S proteasome is essential for the degradation of ubiquitin-modified proteins (Smalle et al., 2004). We then examined SUMOylation and ubiquitination in transgenic plants. As shown in Fig. 7B, both transgenic plants had higher levels of SUMOylation and ubiquitination after drought stress. Compared with that of *MdDREB2A* OE plants, the SUMOylation level of *MdDREB2A*^*K192R*^ OE plants was much lower. However, their ubiquitination level was also lower (Fig. 7B), suggesting that SUMOylated MdDREB2A may undergo ubiquitination in response to drought in *MdDREB2A* OE plants.

**Fig. 7.**
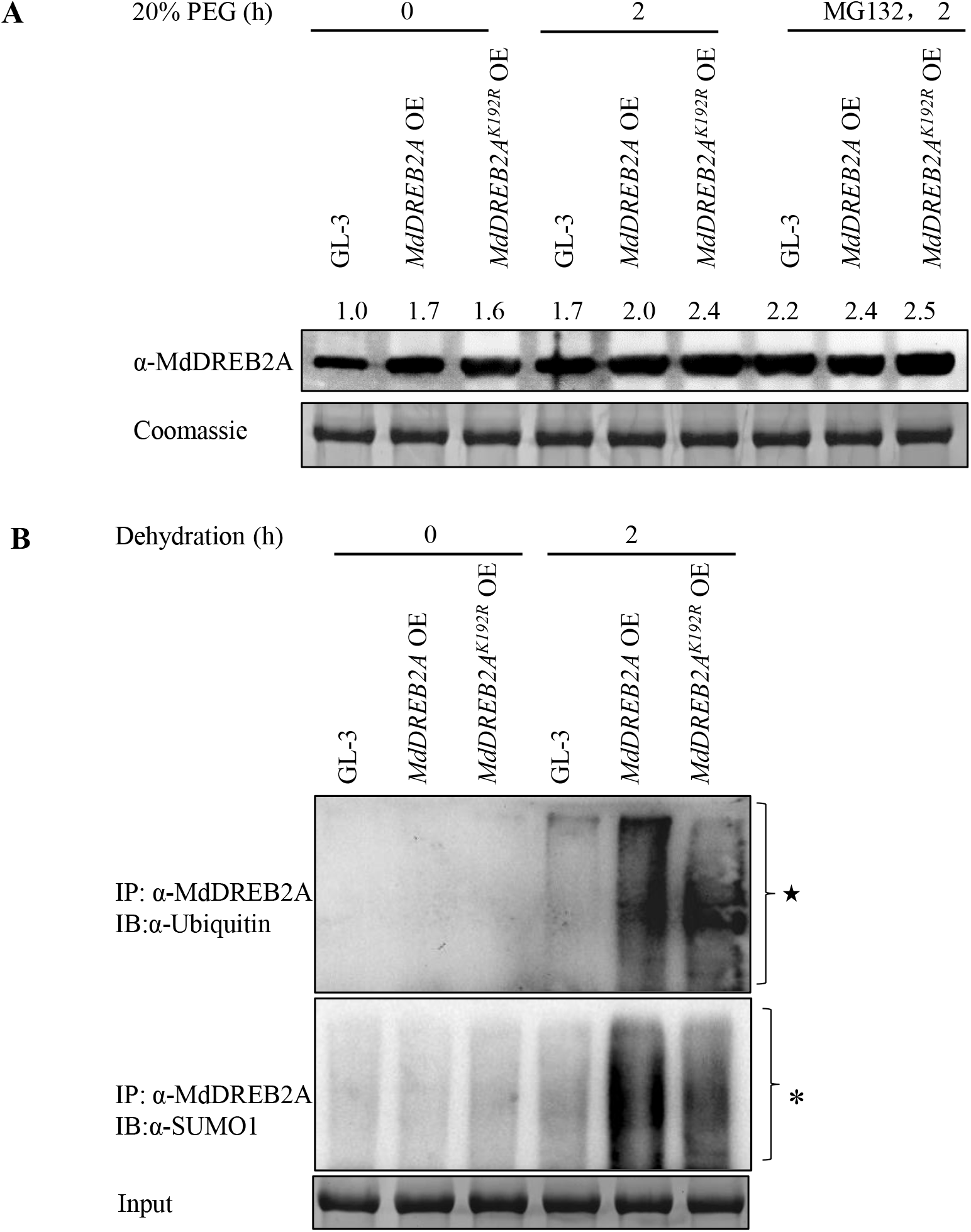
SUMOylation of MdDREB2A couples with ubiquitination mediated by 26S proteasome pathway under drought stress. **(A)** MdDREB2A accumulation in *MdDREB2A* OE and *MdDREB2A*^*K192R*^ OE transgenic plants under simulated drought stress with or without 50 μM MG132 treatment. **(B)** Ubiquitination and SUMOylation of MdDREB2A in *MdDREB2A* OE and *MdDREB2A*^*K192R*^ OE plants in response to dehydration. * indicates SUMOylated substrates; ★ indicates ubiquitinated substrates. OE, overexpression.

### MdRNF4 mediates ubiquitination of SUMOylated MdDREB2A

To identify the proteins responsible for the ubiquitination of SUMOylated MdDREB2A, we performed affinity purified mass spectrometry (AP-MASS) analysis of MdDREB2A under control and drought stress conditions. We identified 1414 and 1472 proteins that may associate with MdDREB2A *in planta* under control and drought conditions, respectively (Supplemental Data set 2). One of the potential MdDREB2A interacting proteins under drought stress was MdRNF4, which encodes an E3 ubiquitin ligase. Homologs of MdRNF4 in mammalian cells and yeast target SUMOylated proteins for degradation by the proteasome pathway (Sun et al., 2007; Tatham et al., 2008; Kumar et al., 2017). We verified the *in vivo* association of MdDREB2A with MdRNF4 using co-immunoprecipitation (Co-IP) analysis (Fig. S18). MdRNF4 contains two SUMO interacting motifs (SIMs) (Fig. S19A). To investigate whether SUMO could be bound to the SIMs of MdRNF4, we performed an Y2H analysis and found that MdRNF4 could interact with MdSUMO2A. When both SIMs were deleted, no interaction was detected. However, deletions of only one SIM did not impair the interaction, indicating that both SIMs are required for the interaction of MdSUMO2A with MdRNF4 (Fig. S19B–C). A microscale thermophoresis (MST) approach and Co-IP assay further verified the interaction between MdSUMO2A and MdRNF4 (Fig. S19D-E).

RNF4 is a SUMO-targeted ubiquitin E3 ligase that is required for degradation of SUMOylated substrates in mammals (Valérie et al., 2008; Geoffroy and Hay, 2009; Zhang et al., 2017a) and the fission yeast *Schizosaccharomyces pombe* (Sun et al., 2007). We hypothesized that this protein is responsible for the ubiquitination of SUMOylated MdDREB2A. To test our hypothesis, we extracted total proteins from GL-3 and *MdDREB2A* OE plants under drought stress and then added purified MdRNF4 to the protein extracts for specific durations. The addition of MdRNF4 for 2 h increased the ubiquitination level of MdDREB2A. However, greater MdDREB2A ubiquitination was observed in *MdDREB2A* OE plants that had higher MdDREB2A SUMOylation levels (Fig. 8A). When MG132 was applied, the ubiquitination of MdDREB2A decreased. To further confirm the requirement of MdRNF4 for degradation of SUMOylated MdDREB2A, we generated transgenic plants with a reduced level of *MdRNF4* (Fig. S20). After immunoprecipitation with anti-MdDREB2A antibody, MdDREB2A ubiquitination decreased in *MdRNF4* RNAi plants under drought conditions (Fig. 8B), further suggesting that MdRNF4 mediates the ubiquitination of MdDREB2A.

**Fig. 8.**
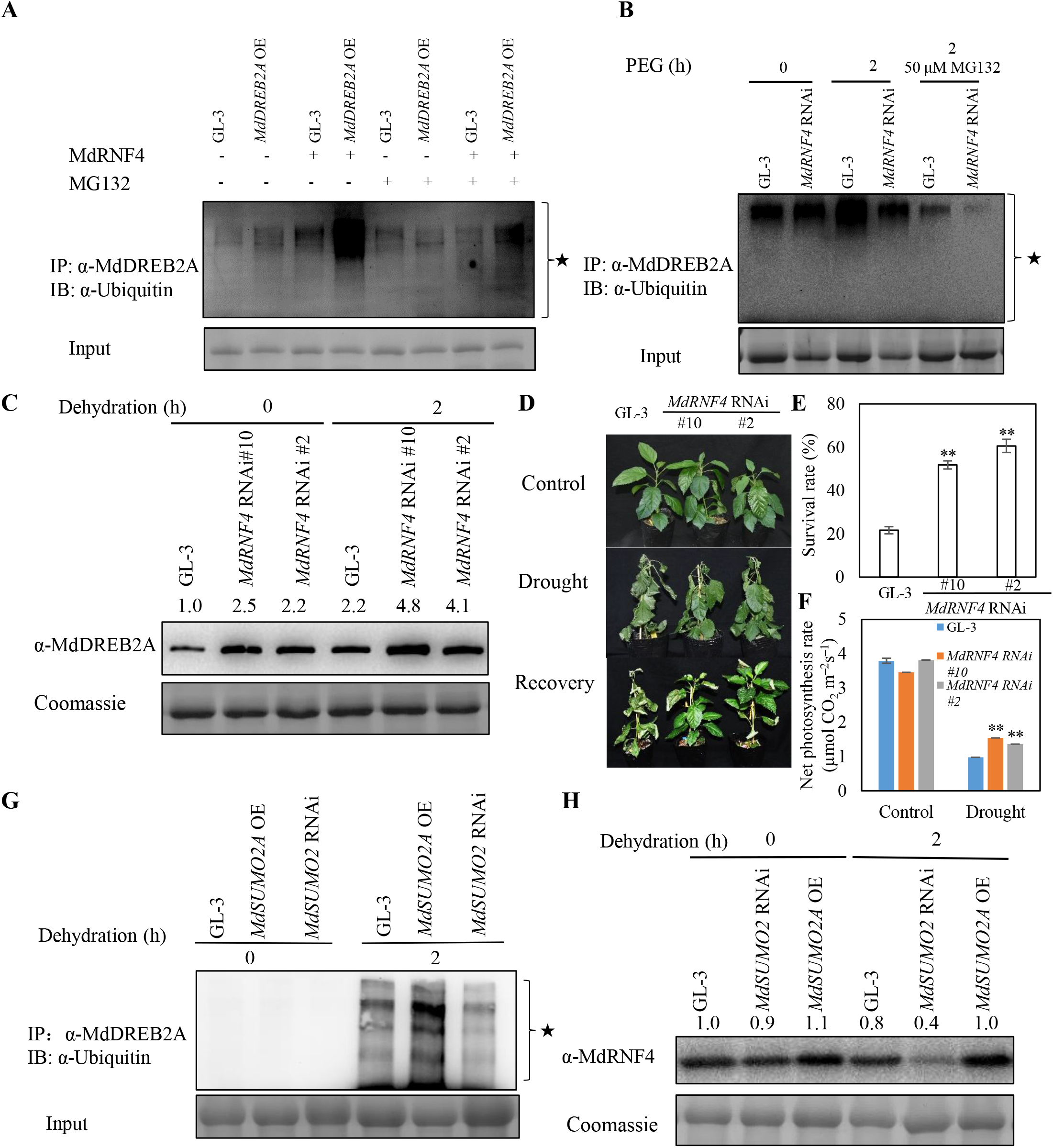
MdRNF4 mediates degradation of SUMOylated MdDREB2A under dehydration conditions. **(A)** Effects of recombinant MdRNF4 on ubiquitination of MdDREB2A under simulated drought stress. Proteins were extracted from PEG-treated GL-3 and *MdDREB2A* OE plants, and recombinant MdRNF4 or 50 μM MG132 was added. **(B)** Ubiquitination of MdDREB2A in *MdRNF4* RNAi plants in response to dehydration. **(C)** MdDREB2A accumulation in *MdRNF4* RNAi plants in response to dehydration. **(D)** Morphology of GL-3 and *MdRNF4* RNAi plants under control and drought stress conditions. **(E)** and **(F)** Survival rate **(E)** and photosynthetic capacity **(F)** of the plants shown in **(D). (G)** Ubiquitination of MdDREB2A in *MdSUMO2* RNAi or *MdSUMO2A* OE plants in response to dehydration. **(H)** MdRNF4 level in *MdSUMO2* RNAi or *MdSUMO2A* OE plants in response to dehydration. Error bars indicate standard error [n = 8 in (D), 3 in (E), 20 in (F)]. Asterisks indicate significant differences based on one-way ANOVA and Tukey test (*, *P* < 0.05; **, *P* < 0.01). OE, overexpression. ★ indicates ubiquitinated substrates. OE, overexpression.

To further analyze the modulation of MdDREB2A stability by MdRNF4, we performed immunoblot analysis of plants under control and drought conditions. MdDREB2A protein levels were higher in *MdRNF4* RNAi plants than in GL-3 plants, either under control or drought conditions; although drought stress induced MdDREB2A accumulation (Fig. 8C). In addition, the *MdRNF4* RNAi plants were more tolerant to drought stress, consistent with the increased tolerance of *MdRNF4* RNAi calli to simulated drought (Fig. 8D-F; Fig. S21A-C).

Because SUMO2s affect MdDREB2A SUMOylation and stability (Fig. 5G), we asked whether this effect was related to MdRNF4. We examined the ubiquitination of MdDREB2A in *MdSUMO2* transgenic plants under control and drought conditions. After drought stress, *MdSUMO2A* OE plants had higher levels of MdDREB2A ubiquitination, and *MdSUMO2* RNAi plants had lower levels (Fig. 8G). In addition, less MdRNF4 accumulated in *MdSUMO2* RNAi plants under drought stress, while more in *MdSUMO2A* OE plants (Fig. 8H), implying the involvement of ubiquitination mediated by MdRNF4 in the MdDREB2A SUMOylation and stability.

## Discussion

Drought stress is one of the major environmental fluctuations that affect plant productivity and survival (Geng et al., 2018; Li et al., 2019). During evolution, plants have acquired divergent strategies to respond to water deficiency, including shortening their life cycles to complete vegetative growth and reproduction before soil water is depleted, evolving unique morphologies and root systems to avoid drought stress, and developing the ability to withstand low tissue water content under drought stress. The latter ability may involve processes such as osmotic adjustment, cellular elasticity, and epicuticular wax formation (Polania et al., 2016; Wei et al., 2016; Yıldırım and Kaya, 2017). In our study, both *MdSUMO2A* OE plants and *MdSUMO2A* RNAi plants were more drought tolerant than the wild type. The *MdSUMO2A* OE plants exhibited greater root system development, more vigorous growth, and higher photosynthetic capacity and hydraulic conductivity (Fig. 2-3; Fig. S4–11). The *MdSUMO2A* RNAi transgenic plants had smaller but thicker leaves, much lower stomatal conductance, and higher water use efficiency (Fig. 2-3; Fig. S4-11). However, the *MdSUMO2A* RNAi plants had a much higher survival rate than the *MdSUMO2A* OE plants. These results suggested that both increased and decreased SUMOylation levels can increase plant drought tolerance.

SUMO is a crucial post-translational modifier in plants that is covalently conjugated with target substrates to maintain chromatin integrity, transduce signals, stabilize proteins, and change cell locations (Dohmen, 2004; Elrouby, 2015; Rytz et al., 2016). Previous studies identified a large number of SUMO substrates in Arabidopsis under heat and oxidative stress, including TPL (TOPLESS), ARF, JAZ, ABF, and NAC proteins (Miller et al., 2010; Rytz et al., 2016; Rytz et al., 2018). Here, we identified 1314 potential targets modified by MdSUMO2A in apple (Supplemental Data set 1). Some MdSUMO2A target proteins were homologous to proteins in Arabidopsis, whereas the majority was unique proteins in the apple genome. The reconstituted Arabidopsis SUMOylation cascade in *E. coli* is a rapid and effective method for evaluating the SUMOylation of potential SUMO target proteins (Okada et al., 2009; Saitoh et al., 2009). We used the apple SUMOylation cascade in *E. coli* as a powerful tool to elucidate the SUMOylation level of targets and confirmed that MdDREB2A, MdALI, and MdAQP2 were MdSUMO2 substrates in apple (Fig. 5), highlighting the power and reliability of this system.

DREB2A encodes a transcription factor that binds to the dehydration-responsive element (DRE) (Yamaguchishinozaki and Shinozaki, 1994; Liu et al., 1998). Numerous studies have reported the positive role of DREB2A in response to drought stress in various plants, including apple, Arabidopsis, rice, maize, and *Pennisetum glaucum*. However, DREB2A sequences from these species did not show high similarity outside of the conserved DNA binding domain in the N-terminal region that may function as a nuclear localization signal (Agarwal et al., 2007; Qin et al., 2007; Qin et al., 2008; Cui et al., 2011; Liao et al., 2016). The N-terminal region of DREB2A that contains the DNA binding and NRD domains is responsible for its protein stability. The DREB2A NRD domain has been shown to interact with DRIP and BPM ubiquitin E3 ligases, leading to ubiquitination and degradation of DREB2A (Qin et al., 2008; Morimoto et al., 2017). DREB2A can be transformed into a stable and constitutively active form (DREB2A-CA) by deleting its NRD domain, thereby facilitating plant drought and heat stress tolerance (Sakuma et al., 2006a). In addition, SUMOylation of DREB2A can increase its protein stability under heat stress by suppressing its interaction with BPM2 (Wang et al., 2020). However, apple MdDREB2A does not contain the NRD domain (Fig. S22). Whether apple MdDREB2A undergoes any protein modifications was previously unknown. In this study, we found that MdDREB2A was a SUMOylation target of the MdSUMO2s (Fig. 5A–C). The critical SUMOylation site of MdDREB2A by MdSUMO2A was the K192 (Fig. 5A). Similar to DREB2As in other plant species, MdDRBE2A was a positive regulator of apple drought stress resistance (Fig. 6A–C). SUMOylation of targets often increases their stability, as well as overall environmental stress resistance (Miura et al., 2007; Zhou et al., 2017; Wang et al., 2020). To our surprise, we found that the mutation of K192 to R caused MdDREB2A protein levels to be more stable in *MdDREB2A*^*K192R*^ OE plants (Fig. 6D). In addition, transgenic plants carrying *MdDREB2A*^*K192R*^ had a higher survival rate than *MdDREB2A* OE plants, implying that DREB2A SUMOylation may serve different functions and proceed by different mechanisms in different plant species in response to stress.

In addition to its covalent attachment to target substrates, SUMO can also interact noncovalently with proteins that contain SIMs (Sun et al., 2007; Nabil et al., 2014; Kumar et al., 2017). SIPs in mammals and Arabidopsis include ubiquitin E3 ligases, DNA methyltransferases or demethylases, and histone methyltransferases or demethylases (Nabil et al., 2014; Kumar et al., 2017). We also identified SIPs in the apple genome and obtained similar results (Supplemental Data set 3). Among the SIPs, we identified a RING finger protein 4, MdRNF4, that appeared with the highest frequency in the Y2H screen. Similar RING-type ubiquitin E3 ligases (RNF4s) have been reported to interact with SUMO and ubiquitinate SUMOylated substrates via the 26S proteasome in mammals and yeast (Sun et al., 2007; Valérie et al., 2008; Geoffroy and Hay, 2009; Zhang et al., 2017a). Our study found that MdRNF4 mediated ubiquitination of SUMOylated MdDREB2A by a 26S proteasome pathway, resulting in the degradation of SUMOylated MdDREB2A (Fig. 8). These results suggest a widely conserved function for RNF4 in ubiquitination among eukaryotes.

In summary, we investigated the relationship between SUMOylation and drought stress tolerance in perennial apple trees. Using *MdSUMO2A* OE and *MdSUMO2* RNAi plants, we observed that both decreased and increased SUMOylation can increase plant drought tolerance, although decreased SUMOylation was associated with relatively higher survival rates. We also showed that increased SUMOylation of MdDREB2A in *MdSUMO2A* OE and *MdDREB2A* OE plants was associated with MdRNF4-mediated greater ubiquitination under drought stress, thereby relatively decreasing MdDREB2A accumulation in *MdSUMO2A* OE and *MdDREB2A* OE plants compared with *MdSUMO2* RNAi and *MdDREB2A*^*K192R*^ OE plants.

## Methods

### Plant materials and growth conditions

The experiments were conducted at Northwest A&F University, Yangling, China (34°20’N, 108°24’E). The transgenic lines and GL-3 plants after rooting on MS were transplanted to soil and grown for 3 months at 25°C under a long day photoperiod (14 h : 10 h, light : dark). The general management was conducted using the method described by Xie (Xie et al., 2017).

The leaves of apple ‘Golden delicious’ (*Malus* x *domestica*) were used for gene cloning. A line isolated from ‘Royal Gala’ (*Malus* x *domestica*) named GL-3 (Dai et al., 2013), which has high regeneration capacity, was used for genetic transformation. GL-3 tissue-cultured plants were subcultured every 4 weeks. They were grown on MS medium (4.43 g/L MS salts, 30 g/L sucrose, 0.2 mg/L 6-BA, 0.2 mg/L IAA, and 7.5 g/L agar, pH 5.8) under long-day conditions (14 h : 10 h, light : dark) at 25°C.

### Generation of transgenic apple plants and calli

201-bp (3’ UTR region), 121-bp (conserved CDS of *MdSUMO2s*), or 74-bp fragments of *MdSUMO2A, MdSUMO2*, or *MdRNF4* were individually cloned into the pDONR222 vector by multisite Gateway recombination, as described by Karimi et al. (Karimi and Hilson, 2007) and subsequently transferred to RNA silencing vector pK7GWIWG2, a destination vector containing an N-terminal GFP tag by LR recombination. To overexpress genes, the coding sequences of *MdSUMO2A, MdDREB2A*, or *MdDREB2A*^*K192R*^ were constructed to pCambia 2300 with N-myc tag or pGWB418. All the constructed vectors were transformed into *Agrobacterium* strain *EHA105. Agrobacterium*-mediated transformation of apple was carried out as described, using GL-3 as the genetic background (Holefors et al., 1998; Dai et al., 2013).

To generate transgenic apple calli, ‘Orin’ (*Malus*× *domestica*) calli grown on MS media (1.5 mg/ L 2,4-Dichlorophenoxyacetic acid (2,4-D), and 0.4 mg/L 6-BA at the dark environment) were used as the wild type. The coding sequence of *MdDREB2A, MdDREB2A*^*K192R*^, or *MdRNF4* was cloned into plant binary vector pGWB418. A 300-bp sequence of *MdRNF4* was cloned into pK7GWIWG2 to knock down *MdRNF4* expression. The resulting plasmids were transformed into *Agrobacterium* strain *EHA105* and then tranformed to ‘Orin’ calli according to previous methods (An et al., 2019; An et al., 2020) The primers used for constructing these vectors are shown in Supplemental Data Set 4.

### Stress treatment

For long-term drought treatment, 3-month-old GL-3 and transgenic apple plants were transplanted to a greenhouse at the beginning of April, 2019. The drought treatment was performed two months later in June. The plants were grown in plastic pots (15 cm × 20 cm, ∼1.3 L) filled with a mixture of sand and substrate (PINDSTRUP, Denmark) (1:1, v/v). The measurement of soil volumetric water content (VWC) was conducted by TDR (FS6430,USA). At the beginning of drought treatment, uniform trees of each line (30 trees for each line) were divided into two groups for the following treatments (15 trees for each treatment for each line): (1) control, well-watered, irrigated daily to maintain 43-48% of VWC and (2) moderate drought, irrigated daily to maintain 18-23% of VWC. The treatment was lasted for three months. The photosynthetic capacity was determined with LI-Cor 6400 portable photosynthesis system (LI-COR, Huntington Beach, CA, USA). Hydraulic conductivity of roots and shoots were conducted by an HPFM (Dynamax, Houston) as described previously (Geng et al., 2018). The thickness of leaves were measured by using tungsten filament scanning electron microscope (JSM-6360LV, Japan) according to the methods described by Liao (Liao et al., 2016) with modifications. For the detection of leaves δ^13^C ‰, mature leaves were colltected. Leaves were oven-dried at 105°C for 0.5 h, and then 70°C for 3 days to dry completely. Dried leaves were ground and filtered through a sieve (80 holes per cm^2^). The δ^13^C ‰ of leaves was determined with an elementary analysis-isotope ratio mass spectrometer (Flash EA 1112 HT-Delta V Advantages, Thermo Fisher Scientific) as described previously (Wang et al., 2018).

For short-term drought treatment, 3-month-old uniform trees of GL-3 and transgenic apples were used. Before treatment, plants were irrigated to maintain saturation of soil water content. Then plants were withheld with water until VWC reached 0, and survival rate was calculated after rewatering for one week. The soil VWC was measured by TDR (FS6430, USA).

### RNA extraction and quantitative real-time RT–PCR

Total RNA from apple leaves was extracted by a CTAB method. DNase I (Fermentas) was used to remove residual genomic DNA. We used total RNA to generate cDNA according to the manufacturer’s instructions by using the RevertAid™ First Strand cDNA synthesis kit (Thermo Scientific, USA). The qRT-PCR was performed in a reaction containing GoTaq^®^ qPCR Master Mix (Promega, USA), cDNA, and primers (described in Supplemental Data Set 4) on an CFX96 real-time PCR detection systems (Bio-Rad, USA). *MdMDH* (malate dehydrogenases) was used as the reference gene.

### Subcellular localization

To generate the constructs for subcellular localization assay, coding region of *MdSUMO2A, MdSUMO2B*, or *MdSUMO2C* was amplified and cloned into pEarleyGate104 vector by BP and LR reactions (Invitrogen), and were then transformed into *Agrobacterium* strain *C58C1*. The empety vector pGWB455 which carries 35S::mCherry was also transformed into *Agrobacterium* strain *C58C1*. The *C58C1* carrying the resulting plasmid, 35S::mCherry, and 35S:p19 (p19 is an RNA silencing repressor protein from *Tomato bushy stunt virus*) was coinfiltrated into tobacco leaves (*Nicotiana benthamiana*). Three days later, the leaf epidermal cells were observed by Nikon A1R/A1 confocal microscope system (Nikon, Tokyo, Japan) for yellow fluorescence observation.

For colocalization of MdSUMO2A with MdDREB2A or MdDREB2A^K192R^, the full length sequence of MdDREB2A or MdDREB2A^K192R^ was cloned into pGWB455 and then transformed into *C58C1*. Mature fragments of *MdSUMO2A, MdSUMO2B* and *MdSUMO2C* (*MdSUMO2*s with exposed GG) were amplified and individually cloned into pEarleyGate104 vector by BP and LR reactions (Invitrogen), and were then transformed into *Agrobacterium* strain *C58C1*. The C58C1 carrying mCherry-MdDREB2A, 35S:p19, and YFP-MdSUMO2A, YFP-MdSUMO2B, or YFP-MdSUMO2C were resuspended in the buffer containing 10 mM MgCl_2_, 10 mM MES–KOH, 180 μM acetosyringone and then co-infiltrated into the tobacco leaves for 3 d to detect signals with confocal microscope. The primers used are listed in Supplemental Data Set 4.

### Histochemical and fluorometric assays for GUS activity

For the promoter-GUS reporter assay, an ∼1000 bp DNA fragment upstream of the *MdSUMO2A* was cloned into pMDC164, and then transformed into *Agrobacterium* strain *GV3101*. The resulting plasmid was introduced into Col-0 using the floral-dipping method (Clough and Bent, 1998) for stable transformation in Arabidopsis. GUS activity was observed after staining with 0.5 mg/mL 5-bromo-4-chloro-3-indolyl-b-Dglucoronide as described previously (Guan et al., 2013). The primers used are listed in Supplemental Data Set 4.

### Endogenous ABA determination

After three months of moderate drought treatment, the mature leaves were collected from GL-3 and *MdSUMO2* transgenic lines to determine ABA content. Leaves were weighed and immediately frozen in liquid nitrogen. Frozen leaves were then pulverized and ABA was extracted as described previously (Chen et al., 2012; Xie et al., 2020). Quantitative determination of endogenous ABA was performed on a UPLC–MS/MS system (QTRAP™ 5500 LC/MS/MS, USA) and a Shimadzu LC-30AD UPLC system (Tokyo, Japan).

### SUMOylation assay in *E. coli*

SUMOylation assays in *E. coli* were conducted as described previously (Elrouby and Coupland, 2010). The coding region of *MdAE1* or *MdAE2* were amplified and cloned into binary expression vector pCDFDuet-1, and mature *MdSUMO2*s or *MdCE* was cloned into pACYCDuet-1. Prokaryotic expression vector PGEX-4T-1 was used to express GST-MdDREB2A, MdALI, and MdAQP2 protein. Subsequently, the resulting plasmids in certain combination were introduced into *Escherichia coli stain* BL21 (DE3). After incubation at 37°C until OD_600_ reached 0.6, 1 mM IPTG (Isopropyl β-D-1-thiogalactopyranoside) was added to induce protein expression. Eight hours later, the bacterium was harvested and denaturized for western blot analysis with GST antibody (M20007, Abmart). The primers used are listed in Supplemental Data Set 4.

### Immunoblot analysis

The proteins of transgenic apple plants and GL-3 were extracted with protein extraction buffer [50 mM Tris-HCl, pH 8.0, 150 mM NaCl, 2 mM EDTA, 1 mM DTT, 10% glycerol, 1% Triton X-100, 1 mM phenylmethylsulfonyl fluoride (PMSF), and 1 × Halt protease inhibitor cocktail (Fisher Scientific)] and centrifuged at 14,000 g at 4°C for 10 min. The extracted proteins were used for western blot analysis with polyclonal MdDREB2A antibody against rabbit, or anti-SUMO (ab5316, Abcam), anti-Ubiquitin (P4D1, Cell Signaling Technology^®^), anti-MdRNF4 (rabbit polyclonal antibody, ABclonal Technology), or anti-Actin (AC009, ABclonal Technology).

### *In vivo* SUMOylation and ubiquitination analysis

Total proteins extracted from transgenic plants (*MdSUMO2A* OE, *MdSUMO2* RNAi, *MdDREB2A* OE, *MdDREB2A*^*K192R*^ OE, *MdRNF4* RNAi, and GL-3) were immunoprecipitated with anti-MdDREB2A and immunoblotted with anti-SUMO (ab5316, Abcam), or anti-Ubiquitin (P4D1, Cell Signaling Technology^®^) antibodies. To examine the ubiquitination and SUMOyltion of MdDREB2A under drought stress conditions, plants were dehydrated for 2 hours.

To dectect the effects of recombinant MdRNF4 on ubiquitination and SUMOylation of MdDREB2A under simulated drought stress, proteins were extracted from PEG-treated GL-3 and *MdDREB2A* OE plants, and recombinant MdRNF4 or 50 μM MG132 was added for 2 hours (An et al., 2019; An et al., 2020). Total proteins were extracted and immunoprecipitated with anti-MdDREB2A and immunoblotted with anti-SUMO (ab5316, Abcam), or anti-Ubiquitin (P4D1, Cell Signaling Technology^®^) antibodies.

### Yeast two-hybrid assay

To identify MdSUMO2 interacting proteins, 1-95 aa of MdSUMO2A (mature MdSUMO2A with exposed GG) was amplified and cloned into pGBKT7 vector to generate bait plasmid. Y2H screen was performed to screen the apple library according to the user manual of Matchmaker™ Gold Yeast Two Hybrid System (Clontech, Japan) by using *Saccharomyces cerevisiae* strain Y2H Gold.

To perform the point-to-point Y2H, full length MdSUMO2A was cloned into pGBKT7, resulting in MdSUMO2A-pGBKT7. Full-length or truncated MdRNF4 with SIM deltion was constructed to pGADT7 vector. MdSUMO2A-pGBKT7 and MdRNF4-pGADT7 or truncated MdRNF4-pGADT7 were co-transformed into yeast strain Y2H Gold. The positive clones were selected on SD-Leu-Trp, and then on SD-Leu-Trp-His-Ade + x-α-gal plates for growth observation and the x-α-gal assay. The primers used are listed in Supplemental Data Set 4.

### CO-IP assay

For Co-IP analysis, the leaves of GL-3 were dehydrated for 2 hours. Total proteins were extracted from leaf samples with extraction buffer [50 mM Tris-HCl, pH 8.0, 150 mM NaCl, 2 mM EDTA, 1 mM DTT, 10% glycerol, 1% Triton X-100, 1 mM phenylmethylsulfonyl fluoride (PMSF), and 1 × Halt protease inhibitor cocktail (Fisher Scientific)]. The protein extracts were incubated overnight with polyclonal MdDREB2A antibody. The immunocomplexes were collected by adding protein A/G agarose beads (Thermo Fisher) and were washed with immunoprecipitation buffer [50 mM Tris-HCl, pH 8.0, 150 mM NaCl, 2 mM EDTA, 1 mM DTT, 10% glycerol, 0.15% Triton X-100, 1 mM PMSF, and 1× Halt protease inhibitor cocktail (Fisher Scientific]]. The pellet (immunocomplexes with beads) was resuspended in 1× SDS-PAGE loading buffer. Eluted proteins were analyzed by immunoblotting using anti-MdRNF4 antibody or anti-MdDREB2A antibody. Chemiluminescence signals were detected by autoradiography.

### AP-MASS assay

To identify the interacting proteins of MdDREB2A *in vivo*, AP-MASS assay was performed as described previously (Maio et al., 2020) with modifications. Total proteins were extracted in leaves of GL-3 plants with or without 2 h dehydation treatments using extraction buffer [50 mM Tris-HCl, pH 8.0, 150 mM NaCl, 2 mM EDTA, 1 mM DTT, 10% glycerol, 1% Triton X-100, 1 mM phenylmethylsulfonyl fluoride (PMSF), and 1× Halt protease inhibitor cocktail (Fisher Scientific)]. The protein extracts were incubated overnight with polyclonal MdDREB2A antibody and then added protein A/G agarose beads (Thermo Fisher) to incubate at 4 °C for additional 4-5 hours. After incubation, the beads were captured with a magnetic rack and washed three times in 0.5 ml of washing buffer (10 mM Tris–HCl pH 7.5, 150 mM NaCl, 0.5 mM EDTA, 1 mM PMSF protease inhibitor). The pellet (immunocomplexes with beads) was resuspended in 1× SDS-PAGE loading buffer and subjected to mass spectrometry analysis (Applied protein technology, China).

### Microscale thermophoresis (MST) assay

Full length of MdSUMO2A and MdDREB2A were cloned into pET-32a. MdCE or MdRNF4 was cloned into pGEX-4T and pMAL-c5X, respectively. The resulting plasmids were expressed in *E. coli* BL21. Recombinant protein MdSUMO2A-HIS and MdDREB2A were purified by HIS Sepharose beads (GE Healthcare, Fairfield, CT, USA), GST-MdCE was purfied by Pierce™ Glutathione Spin Columns (16105, Thermo Scientific™, USA) and MBP-MdRNF4 was purified by MBP TRAP HP (GE Healthcare). MST was conducted according the manuferturer’s manual (NanoTemper, Germany). The primers used are listed in Supplemental Data Set 4.

### Accession numbers

The accession numbers in GDR are as follows: MdSUMO2B (MD17G1103900, MD09G1113800), MdSUMO2A (MD03G1194700, MD11G1211000), MdSUMO2C (MD05G1173700, MD10G1161600); and in NCBI under the following: MdDREB2A (NP_001280947.1), MdAE1 (XP_028948277.1), MdAE2 (XP_008382303.1), MdCE (XP_008338336.1), MdRNF4 (XP_008346210.1), MdALI (XP_008341016.1), MdAQP2 (XP_008363507.1).

## Acknowledgements

This work was supported by the National Key Research and Development Program of China (2019YFD1000100, 2018 YFD1000100), and National Natural Science Foundation of China (31872080).

## Author contributions

We thank Dr. Zhihong Zhang from Shenyang Agricultural University for providing tissue-cultured GL-3 plants. Q.G. and F.M. designed the project. X.L., S.Z., L.L., H.D., Z.L., P. C., Z.M., S.Z., and B.C. performed the experiments. Q.G., X.L., H.D., B.C. and L.L. analyzed the data. Q.G., X.L. and F.M. wrote the manuscript.

## References

Agarwal, P., Agarwal, P.K., Nair, S., Sopory, S.K., and Reddy, M.K. (2007). Stress-inducible DREB2A transcription factor from Pennisetum glaucum is a phosphoprotein and its phosphorylation negatively regulates its DNA-binding activity. Mol Genet Genomics 277, 189.

An, J.P., Wang, X.F., Zhang, X.W., Bi, S.Q., You, C.X., and Hao, Y.J. (2019). MdBBX22 regulates UV-B-induced anthocyanin biosynthesis through regulating the function of MdHY5 and is targeted by MdBT2 for 26S proteasome-mediated degradation. Plant Biotechnol J 17, 2231–2233.

An, J.P., Wang, X.F., Espley, R.V., Lin-Wang, K., Bi, S.Q., You, C.X., and Hao, Y.J. (2020). An apple B-Box protein MdBBX37 modulates anthocyanin biosynthesis and hypocotyl elongation synergistically with MdMYBs and MdHY5. Plant Cell Physiol. 61, 130–143.

Anyia, A.O., and Herzog, H. (2004). Water-use efficiency, leaf area and leaf gas exchange of cowpeas under mid-season drought. Europ. J. Agronomy 20, 327–339.

Basu, S., Ramegowda, V., Kumar, A., and Pereira, A. (2016). Plant adaptation to drought stress. F1000research 5, 1554.

Castro, P.H., Tavares, R.M., Bejarano, E.R., and Azevedo, H. (2012). SUMO, a heavyweight player in plant abiotic stress responses. Cell Mol Life Sci 69, 3269–3283.

Catala, R., Ouyang, J., Abreu, I.A., Hu, Y., Seo, H., Zhang, X., and Chua, N.H. (2007). The Arabidopsis E3 SUMO ligase SIZ1 regulates plant growth and drought responses. Plant cell 19, 2952–2966.

Chen, M., Wang, Q., Cheng, X., Xu, Z., Li, L., Ye, X., Xia, L., and Ma, Y. (2007). GmDREB2, a soybean DRE-binding transcription factor, conferred drought and high-salt tolerance in transgenic plants. Biochem. Biophys. Res. Commun. 353, 299–305.

Chen, M.L., Fu, X.M., Liu, J.Q., Ye, T.T., Hou, S.Y., Huang, Y.Q., Yuan, B.F., Wu, Y., and Feng, Y.Q. (2012). Highly sensitive and quantitative profiling of acidic phytohormones using derivatization approach coupled with nano-LC-ESI-Q-TOF-MS analysis. J Chromatogr B Analyt Technol Biomed Life Sci 905, 67–74.

Clough, S., and Bent, A. (1998). Floral dip: a simplified method for Agrobacterium-mediated transformation of Arabidopsis thaliana. Plant J. 16, 735–743.

Colby, T., Matthai, A., Boeckelmann, A., and Stuible, H.P. (2006). SUMO-conjugating and SUMO-deconjugating enzymes from Arabidopsis. Plant Physiol 142, 318–332.

Cui, M., Zhang, W., Zhang, Q., Xu, Z., Zhu, Z., Duan, F., and Wu, R. (2011). Induced over-expression of the transcription factor OsDREB2A improves drought tolerance in rice. Plant physiology and biochemistry : PPB 49, 1384–1391.

Dai, A.G. (2013). Increasing drought under global warming in observations and models. Nature Climate Change 3, 52–58.

Dai, H., Li, W., Han, G., Yang, Y., Ma, Y., Li, H., and Zhang, Z. (2013). Development of a seedling clone with high regeneration capacity and susceptibility to Agrobacterium in apple. Sci. Hortic 164, 202–208.

Dohmen, R.J. (2004). SUMO protein modification. Biochim Biophys Acta 1695, 113–131.

Elrouby, N. (2015). Analysis of small ubiquitin-like modifier (SUMO) targets reflects the essential nature of protein SUMOylation and provides insight to elucidate the role of SUMO in plant development. Plant Physiol 169, 1006–1017.

Elrouby, N., and Coupland, G. (2010). Proteome-wide screens for small ubiquitin-like modifier (SUMO) substrates identify Arabidopsis proteins implicated in diverse biological processes. P Natl Acad Sci USA 107, 17415–17420.

Galanty, Y., Belotserkovskaya, R., Coates, J., and Jackson, S.P. (2012). RNF4, a SUMO-targeted ubiquitin E3 ligase, promotes DNA double-strand break repair. Genes Dev 26, 1179–1195.

Geng, D.L., Shen, X., Xie, Y., Yang, Y., Bian, R., Gao, Y., Li, P., Sun, L., Feng, H., Ma, F., and Guan, Q. (2020). Regulation of phenylpropanoid biosynthesis by MdMYB88 and MdMYB124 contributes to pathogen and drought resistance in apple. Hortic Res 7.

Geng, D.L., Chen, P.X., Shen, X.X., Zhang, Y., Li, X.W., Jiang, L.J., Xie, Y.P., Niu, C.D., Zhang, J., Huang, X.H., Ma, F.W., and Guan, Q.M. (2018). MdMYB88 and MdMYB124 enhance drought tolerance by modulating root vessels and cell walls in apple. Plant Physiol 178, 1296–1309.

Geoffroy, M.-C., and Hay, R.T. (2009). An additional role for SUMO in ubiquitin-mediated proteolysis. Nat Rev Mol Cell Biol 10, 564.

Guan, Q.M., Lu, X.Y., Zeng, H.T., Zhang, Y.Y., and Zhu, J.H. (2013). Heat stress induction of miR398 triggers a regulatory loop that is critical for thermotolerance in Arabidopsis. Plant J 74, 840–851.

Holefors, A., Xue, Z.T., and Welander, M. (1998). Transformation of the apple rootstock M26 with the rolA gene and its influence on growth. J Plant Physiol 136, 69–78.

Hu, L., Xie, Y., Fan, S., Wang, Z., Wang, F., Zhang, B., Li, H., Song, J., and Kong, L. (2018). Comparative analysis of root transcriptome profiles between drought-tolerant and susceptible wheat genotypes in response to water stress. Plant Sci 272, 276.

Karan, R., and Subudhi, P.K. (2012). A stress inducible SUMO conjugating enzyme gene (SaSce9) from a grass halophyte Spartina alterniflora enhances salinity and drought stress tolerance in Arabidopsis. BMC Plant Biol 12, 187–187.

Karimi, M., and Hilson, D.P. (2007). Recombinational cloning with plant gateway vectors. Plant Physiol 145, 1144–1154.

Kim, J.-S., Mizoi, J., Yoshida, T., Fujita, Y., Nakajima, J., Ohori, T., Todaka, D., Nakashima, K., Hirayama, T., Shinozaki, K., and Yamaguchi-Shinozaki, K. (2011). An ABRE promoter sequence is involved in osmotic stress-responsive expression of the DREB2A gene, which encodes a transcription factor regulating drought-inducible genes in Arabidopsis. Plant Cell Physiol 52, 2136–2146.

Kim, J., Song, J., and Seo, H. (2017). Post-translational modifications of Arabidopsis E3 SUMO ligase AtSIZ1 are controlled by environmental conditions. FEBS Open Bio 7, 1622–1634.

Kumar, R., Gonzalez-Prieto, R., Xiao, Z., Verlaan-de Vries, M., and Vertegaal, A.C.O. (2017). The STUbL RNF4 regulates protein group SUMOylation by targeting the SUMO conjugation machinery. Nat Commun 8.

Li, X.W., Chen, P.X., Xie, Y.P., Yan, Y., Wang, L.P., Dang, H., Zhang, J., Xu, L.Y., Ma, F.W., and Guan, Q.M. (2020). Apple SERRATE negatively mediates drought resistance by regulating MdMYB88 and MdMYB124 and microRNA biogenesis. Hortic Res 7.

Li, X.W., Xie, Y., Lu, L., Yan, M., Fang, N., Xu, J., Wang, L., Yan, Y., Zhao, T., van Nocker, S., Ma, F., Liang, D., and Guan, Q. (2019). Contribution of methylation regulation of MpDREB2A promoter to drought resistance of Mauls prunifolia. Plant Soil 441, 15–32.

Liao, X., Guo, X., Wang, Q., Wang, Y., Zhao, D., Yao, L., Wang, S., Liu, G., and Li, T. (2016). Overexpression of MsDREB6. 2 results in cytokinin-deficient developmental phenotypes and enhances drought tolerance in transgenic apple plants. Plant J 89.

Liu, Q., Kasuga, M., Sakuma, Y., Abe, H., Miura, S., Yamaguchi-Shinozaki, K., and Shinozaki, K. (1998). Two transcription factors, DREB1 and DREB2, with an EREBP/AP2 DNA binding domain separate two cellular signal transduction pathways in drought-and low-temperature-responsive gene expression, respectively, in Arabidopsis. Plant Cell 10, 1391–1406.

Maio, F., Helderman, T.A., Arroyo-Mateos, M., van der Wolf, M., Boeren, S., Prins, M., and van den Burg, H.A. (2020). Identification of tomato proteins that interact with replication initiator protein (Rep) of the geminivirus TYLCV. Front Plant Sci 11, 1069.

Mao, H., Wang, H., Liu, S., Li, Z., Yang, X., Yan, J., Li, J., Tran, L.S., and Qin, F. (2015). A transposable element in a NAC gene is associated with drought tolerance in maize seedlings. Nat Commun 6, 8326.

Meng, C., Zhang, W., Zhang, Q., Xu, Z., Zhu, Z., Duan, F., and Wu, R. (2011). Induced over-expression of the transcription factor OsDREB2A improves drought tolerance in rice. Plant physiology and biochemistry : PPB 49, 1384–1391.

Miller, M.J., and Vierstra, R.D. (2011). Mass spectrometric identification of SUMO substrates provides insights into heat stress-induced SUMOylation in plants. Plant Signal Behav 6, 130–133.

Miller, M.J., Barrett-Wilt, G.A., Hua, Z., and Vierstra, R.D. (2010). Proteomic analyses identify a diverse array of nuclear processes affected by small ubiquitin-like modifier conjugation in Arabidopsis. P Natl Acad Sci USA 107, 16512–16517.

Miura, K., Okamoto, H., Okuma, E., Shiba, H., Kamada, H., Hasegawa, P., and Murata, Y. (2013). SIZ1 deficiency causes reduced stomatal aperture and enhanced drought tolerance via controlling salicylic acid-induced accumulation of reactive oxygen species in Arabidopsis. Plant J 73, 91–104.

Miura, K., Jin, J.B., Lee, J., Yoo, C.Y., Stirm, V., Miura, T., Ashworth, E.N., Bressan, R.A., Yun, D.J., and Hasegawa, P.M. (2007). SIZ1-mediated sumoylation of ICE1 controls CBF3/DREB1A expression and freezing tolerance in Arabidopsis. Plant Cell 19, 1403–1414.

Mizoi, J., Ohori, T., Moriwaki, T., Kidokoro, S., Todaka, D., Maruyama, K., Kusakabe, K., Osakabe, Y., Shinozaki, K., and Yamaguchi-Shinozaki, K. (2013). GmDREB2A;2, a canonical DEHYDRATION-RESPONSIVE ELEMENT-BINDING PROTEIN2-type transcription factor in soybean, is posttranslationally regulated and mediates dehydration-responsive element-dependent gene expression. Plant Physiol 161, 346–361.

Morimoto, K., Ohama, N., Kidokoro, S., Mizoi, J., Takahashi, F., Todaka, D., Mogami, J., Sato, H., Qin, F., Kim, J.-S., Fukao, Y., Fujiwara, M., Shinozaki, K., and Yamaguchi-Shinozaki, K. (2017). BPM-CUL3 E3 ligase modulates thermotolerance by facilitating negative regulatory domain-mediated degradation of DREB2A in Arabidopsis. P Natl Acad Sci USA 114, E8528–E8536.

Nabil, E., Mitzi Villajuana, B., Aimone, P., and George, C. (2014). Identification of Arabidopsis SUMO-interacting proteins that regulate chromatin activity and developmental transitions. P Natl Acad Sci USA 110, 19956–19961.

Neelam, M., Li, S., Xunlu, Z., Jennifer, S., Anurag, P.S., Xiaojie, Y., Necla, P., Nardana, E., Hong, L., and Guoxin, S. (2017). Overexpression of the rice SUMO E3 ligase gene OsSIZ1 in cotton enhances drought and heat tolerance, and substantially improves fiber yields in the field under reduced irrigation and rainfed conditions. Plant Cell Physiol., 735–746.

Niu, C.D., Li, H., Jiang, L., Yan, M., Li, C., Geng, D., Xie, Y., Yan, Y., Shen, X., Chen, P., Dong, J., Ma, F., and Guan, Q. (2019). Genome-wide identification of drought-responsive microRNAs in two sets of Malus from interspecific hybrid progenies. Hortic Res 6, 75.

Novatchkova, M., Budhiraja, R., Coupland, G., Eisenhaber, F., and Bachmair, A. (2004). SUMO conjugation in plants. Planta 220, 1–8.

Nurdiani, D., Widyajayantie, D., and Nugroho, S. (2018). OsSCE1 encoding SUMO E2-conjugating enzyme involves in drought stress response of Oryza sativa. Rice Science 25, 73–81.

Okada, S., Nagabuchi, M., Takamura, Y., Nakagawa, T., Shinmyozu, K., Nakayama, J.-i., and Tanaka, K. (2009). Reconstitution of Arabidopsis thaliana SUMO pathways in E. coli: functional evaluation of SUMO machinery proteins and mapping of SUMOylation sites by mass spectrometry. Plant Cell Physiol 50, 1049–1061.

Polania, J.A., Poschenrieder, C., Beebe, S., and Rao, I.M. (2016). Effective use of water and increased dry matter partitioned to grain contribute to yield of common bean improved for drought resistance. Front Plant Sci 7.

Qin, F., Kakimoto, M., Sakuma, Y., Maruyama, K., Osakabe, Y., Tran, L., Shinozaki, K., and Yamaguchi-Shinozaki, K. (2007). Regulation and functional analysis of ZmDREB2A in response to drought and heat stresses in Zea mays L. Plant J 50, 54–69.

Qin, F., Sakuma, Y., Tran, L.S., Maruyama, K., Kidokoro, S., Fujita, Y., Fujita, M., Umezawa, T., Sawano, Y., Miyazono, K., Tanokura, M., Shinozaki, K., and Yamaguchi-Shinozaki, K. (2008). Arabidopsis DREB2A-interacting proteins function as RING E3 ligases and negatively regulate plant drought stress-responsive gene expression. Plant Cell 20, 1693–1707.

Reis, R.R., Dias Brito da Cunha, B.A., Martins, P.K., Bazzo Martins, M.T., Alekcevetch, J.C., Chalfun-Junior, A., Andrade, A.C., Ribeiro, A.P., Qin, F., Mizoi, J., Yamaguchi-Shinozaki, K., Nakashima, K., Correa Carvalho, J.d.F., Ferreira de Sousa, C.A., Nepomuceno, A.L., Kobayashi, A.K., and Correa Molinari, H.B. (2014). Induced over-expression of ArDREB2A CA improves drought tolerance in sugarcane. Plant Sci 221, 59–68.

Rytz, T., Miller, M., McLoughlin, F., Augustine, R., Marshall, R., Juan, Y., Charng, Y., Scalf, M., Smith, L., and Vierstra, R. (2018). SUMOylome profiling reveals a diverse array of nuclear targets modified by the SUMO ligase SIZ1 during heat stress. Plant Cell 30, 1077–1099.

Rytz, T.C., Miller, M.J., and Vierstra, R.D. (2016). Purification of SUMO Conjugates from Arabidopsis for Mass Spectrometry Analysis. Methods Mol Biol, 257–281.

Sadhukhan, A., Panda, S.K., and Sahoo, L. (2014). The cowpea RING ubiquitin ligase VuDRIP interacts with transcription factor VuDREB2A for regulating abiotic stress responses. Plant Physiol Biochem. 83, 51–56.

Saitoh, H., Uwada, J., and Azusa, K. (2009). Strategies for the expression of SUMO-modified target proteins in Escherichia coli. Methods Mol Biol 497, 211–221.

Sakuma, Y., Maruyama, K., Qin, F., Osakabe, Y., Shinozaki, K., and Yamaguchi-Shinozaki, K. (2006a). Dual function of an Arabidopsis transcription factor DREB2A in water-stress-responsive and heat-stress-responsive gene expression. P Natl Acad Sci USA 103, 18822–18827.

Sakuma, Y., Maruyama, K., Osakabe, Y., Qin, F., Seki, M., Shinozaki, K., and Yamaguchi-Shinozaki, K. (2006b). Functional analysis of an Arabidopsis transcription factor, DREB2A, involved in drought-responsive gene expression. Plant Cell 18, 1292–1309.

Seiler, C., Harshavardhan, V.T., Rajesh, K., Reddy, P.S., Strickert, M., Rolletschek, H., Scholz, U., Wobus, U., and Sreenivasulu, N. (2011). ABA biosynthesis and degradation contributing to ABA homeostasis during barley seed development under control and terminal drought-stress conditions. J Exp Bot 62, 2615–2632.

Smalle, Jan, Vierstra, Richard, and D. (2004). The ubiquitin 26S proteasome proteolytic pathway. Annu Rev Plant Biol 55, 555–590.

Sobko, A., Ma, H., and Firtel, R.A. Regulated SUMOylation and ubiquitination of DdMEK1 is required for proper chemotaxis. Dev Cell 2, 0–756.

Srivastava, A., Zhang, C., Caine, R., Gray, J., and Sadanandom, A. (2017). Rice SUMO protease Overly Tolerant to Salt 1 targets the transcription factor, OsbZIP23 to promote drought tolerance in rice. Plant J.

Sun, H., Leverson, J.D., and Hunter, T. (2007). Conserved function of RNF4 family proteins in eukaryotes: targeting a ubiquitin ligase to SUMOylated proteins. EMBO J 26, 4102–4112.

Sun, J., Gu, J., Zeng, J., Han, S., Song, A., Chen, F., Fang, W., Jiang, J., and Chen, S. (2013a). Changes in leaf morphology, antioxidant activity and photosynthesis capacity in two different drought-tolerant cultivars of chrysanthemum during and after water stress. Sci. Hortic 161, 249–258.

Sun, X., Wang, P., Jia, X., Huo, L.Q., Che, R.M., and Ma, F.W. (2018). Improvement of drought tolerance by overexpressing MdATG18a is mediated by modified antioxidant system and activated autophagy in transgenic apple. Plant Biotechnol J 16, 545–557.

Sun, X.P., Yan, H.L., Kang, X.Y., and Ma, F.W. (2013b). Growth, gas exchange, and water-use efficiency response of two young apple cultivars to drought stress in two scion-one rootstock grafting system. Photosynthetica 51, 404–410.

Tatham, M.H., Marie-Claude, G., Linnan, S., Anna, P., Neil, H., Jaffray, E.G., Palvimo, J.J., and Hay, R.T. (2008). RNF4 is a poly-SUMO-specific E3 ubiquitin ligase required for arsenic-induced PML degradation. Nat Cell Biol 10, 538–546.

Valérie, L.B., Marion, J., Shirine, B., Rihab, N., Ming, L., Laurent, P., Jun, Z., Jun, Z., Brian, R., and Hugues, D.T. (2008). Arsenic degrades PML or PML-RARalpha through a SUMO-triggered RNF4/ubiquitin-mediated pathway. Nat Cell Biol 10, 547–555.

Vierstra, and R., D. (2012). The Expanding Universe of Ubiquitin and Ubiquitin-Like Modifiers. Plant Physiol 160, 2.

Virlet, N., Costes, E., Martinez, S., Kelner, J.J., and Regnard, J.L. (2015). Multispectral airborne imagery in the field reveals genetic determinisms of morphological and transpiration traits of an apple tree hybrid population in response to water deficit. J Exp Bot 66, 5453–5465.

Wang, F., Liu, Y., Shi, Y., Han, D., Wu, Y., Ye, W., Yang, H., Li, G., Cui, F., Wan, S., Lai, J., and Yang, C. (2020). SUMOylation stabilizes the transcription factor DREB2A to improve plant thermotolerance. Plant Physiol 183, 41–50.

Wang, H., Zhao, S., Mao, K., Dong, Q., Liang, B., Li, C., Wei, Z., Li, M., and Ma, F. (2018). Mapping QTLs for water-use efficiency reveals the potential candidate genes involved in regulating the trait in apple under drought stress. BMC Plant Biol 18, 136.

Wei, W., Hu, Y., Han, Y.-T., Zhang, K., Zhao, F.-L., and Feng, J.-Y. (2016). The WRKY transcription factors in the diploid woodland strawberry Fragaria vesca: Identification and expression analysis under biotic and abiotic stresses. Plant Physiol Biochem. 105, 129–144.

Wu, S., Liang, D., and Ma, F. (2014). Leaf micromorphology and sugar may contribute to differences in drought tolerance for two apple cultivars. Plant Physiol Biochem. 80, 249–258.

Xie, Y.P., Bao, C.N., Chen, P.X., Cao, F.G., Liu, X.F., Geng, D.L., Li, Z.X., Li, X.W., Hou, N., Zhi, F., Niu, C.D., Zhou, S.X., Zhan, X.Q., Ma, F.W., and Guan, Q.M. (2020). ABA homeostasis is mediated by a feedback regulation of MdMYB88 and MdMYB124. J Exp Bot. eraa449.

Xie, Y.P., Chen, P.X., Yan, Y., Bao, C.N., Li, X.W., Wang, L.P., Shen, X.X., Li, H., Liu, X.F., Niu, C.D., Zhu, C., Fang, N., Shao, Y., Zhao, T., Yu, J., Zhu, J., Xu, L., van Nocker, S., Ma, F.W., and Guan, Q.M. (2017). An atypical R2R3 MYB transcription factor increases cold hardiness by CBF-dependent and CBF-independent pathways in apple. New Phytol 218, 201–218.

Xu, G., Duan, B., and Li, C. (2008). Different adaptive responses of leaf physiological and biochemical aspects to drought in two contrasting populations of seabuckthorn. Can. J. For. Res. 38, 584–591.

Yamaguchishinozaki, K., and Shinozaki, K. (1994). A novel cis-acting element in an Arabidopsis gene is involved in responsiveness to drought, low-temperature, or high-salt stress. Plant Cell 6, 251–264.

Yıldırım, K., and Kaya, Z. (2017). Gene regulation network behind drought escape, avoidance and tolerance strategies in black poplar (Populus nigra L.). Plant Physiol Biochem 115, 183–199.

Yordanov, I., Velikova, V., and Tsonev, T. (2000). Plant responses to drought, acclimation, and stress tolerance. Photosynthetica 38, 171–186.

Zhang, L., Xie, F., Zhang, J., Dijke, P.T., and Zhou, F. (2017a). SUMO-triggered ubiquitination of NR4A1 controls macrophage cell death. Cell Death & Differentiation 24.

Zhang, S., Qi, Y., Liu, M., and Yang, C. (2013). SUMO E3 ligase AtMMS21 regulates drought tolerance in Arabidopsis thaliana. J Integr Plant Biol 55, 83–95.

Zhang, S., Zhuang, K., Wang, S., Lv, J., Ma, N.N., and Meng, Q. (2017b). A novel tomato SUMO E3 ligase, SlSIZ1, confers drought tolerance in transgenic tobacco. J Integr Plant Biol.

Zhou, L.J., Li, Y.Y., Zhang, R.F., Zhang, C.L., Xie, X.B., Zhao, C., and Hao, Y.J. (2017). The small ubiquitin-like modifier E3 ligase MdSIZ1 promotes anthocyanin accumulation by sumoylating MdMYB1 under low-temperature conditions in apple. Plant Cell Environ 40, 2068–2080.

Zhu, J.K. (2016). Abiotic stress signaling and responses in plants. Cell 167, 313–324.

